# Intergenerational Stress Transmission is Associated with Brain Metabotranscriptome Remodeling and Mitochondrial Dysfunction

**DOI:** 10.1101/2021.04.09.438868

**Authors:** Sammy Alhassen, Siwei Chen, Lamees Alhassen, Alvin Phan, Mohammad Khoudari, Angele De Silva, Huda Barhoosh, Zitong Wang, Chelsea Parrocha, Emily Shapiro, Charity Henrich, Zicheng Wang, Leon Mutesa, Pierre Baldi, Geoffrey W. Abbott, Amal Alachkar

**Affiliations:** Department of Pharmaceutical Sciences, University of California-Irvine, CA 92697; Department of Computer Science, School of Information and Computer Sciences, University of California-Irvine, CA 92697; Institute for Genomics and Bioinformatics, School of Information and Computer Sciences, University of California-Irvine, CA 92697; Bioelectricity Laboratory, Department of Physiology and Biophysics, School of Medicine, University of California-Irvine, CA 92697; Center for Human Genetics, College of Medicine and Health Sciences, University of Rwanda, Kigali

**Author notes:** These authors contributed equally to this work (Sammy Alhassen, Siwei Chen). Corresponding Author Dr. Amal Alachkar Department of Pharmaceutical Sciences University of California, Irvine, CA 92697.

**Keywords:** stress, Prenatal, early-life, two-hit, mitochondria, epigenetics, 2-hydroxyglurate

## Abstract

Intergenerational stress increases lifetime susceptibility to depression and other psychiatric disorders. Whether intergenerational stress transmission is a consequence of *in utero* neurodevelopmental disruptions vs early-life mother-infant interaction is largely unknown. Here, we demonstrated that exposure to traumatic stress in mice during pregnancy, through predator scent exposure, induces in the offspring social deficits and depressive-like behavior. We found, through cross-fostering experiments, that raising of normal pups by traumatized mothers produced a similar behavioral phenotype to that induced in pups raised by their biological traumatized mothers. Good caregiving (by non-traumatized mothers), however, did not completely protect against the prenatal trauma-induced behavioral deficits. These findings support a two-hit stress mechanism of both *in utero* and early-life parenting (poor caregiving by the traumatized mothers) environments. Associated with the behavioral deficits, we found profound changes in brain metabolomics and transcriptomic (metabotranscriptome). Striking increases in the mitochondrial hypoxia marker and epigenetic modifier 2-hydroxyglutaric acid, in the brains of neonatal and adult pups whose mothers were exposed to stress during pregnancy, indicated mitochondrial metabolism dysfunctions and epigenetic mechanisms. Bioinformatic analyses revealed mechanisms involving stress- and hypoxia-response metabolic pathways in the brains of the neonatal mice, which appear to lead to long-lasting alterations in mitochondrial-energy metabolism, and epigenetic processes pertaining to DNA and chromatin modifications. Most strikingly, we demonstrated that an early pharmacological intervention that can correct mitochondria metabolism - lipid metabolism and epigenetic modifications with acetyl-L-carnitine (ALCAR) supplementation - produces long-lasting protection against the behavioral deficits associated with intergenerational transmission of traumatic stress.

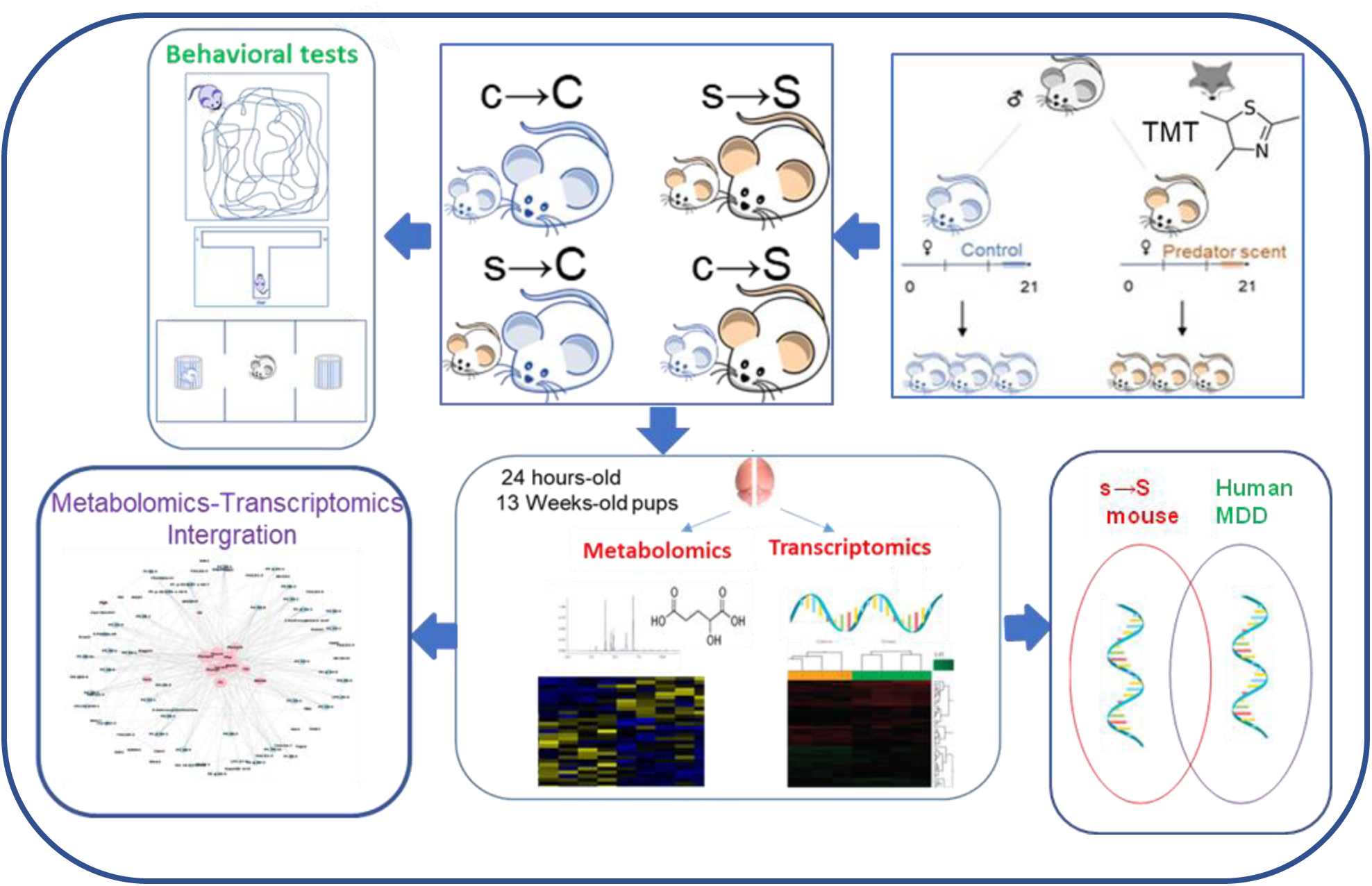

## Introduction

Intergenerational trauma increases lifetime susceptibility to depression and is a major risk factor for developing multiple neuropsychiatric disorders such as post-traumatic stress disorder (PTSD), autism spectrum disorder (ASD), and schizophrenia [1–4]. Depression disorders affect millions of people globally who have been subject to stress and trauma at some times of their lives. While stress deleteriously impacts humans worldwide and attacks all segments of societies at this moment in human history, it is disproportionately devastating to individuals in vulnerable situations such as pregnant women. Unlike other vulnerable conditions, the deleterious impacts of stress during pregnancy are doubled as they affect the mental health of the mothers and the unborn offspring. Human natural experiments provide evidence for the devastating health consequences in offspring as a result of exposure during pregnancy to existential and acute trauma such as war and natural disasters [5–7].

Whether intergenerational trauma transmission and its negative outcomes are a consequence of *in utero* fetal neurodevelopment disruptions or from poor maternal care by traumatized mothers is still largely ambiguous. This ambiguity is mainly due to the complexity, superposition, and inseparability of prenatal and postnatal mechanisms that are not mutually exclusive.

Converging evidence from human and animal studies supports that exposure to stress during pregnancy negatively affects maternal behavior towards, and the care of, offspring [8–10]. On the other hand, maternal stress can affect offspring mental outcomes through prenatal programming, particularly epigenetic mechanisms causing permanent neuronal impairments and poor offspring outcomes that persist throughout life [11–13]. Yet, it is difficult to test the mechanistic involvement of prenatal stress independent of stress effects on the mother care. The limitations in our understanding of the neurobiological and pathophysiological mechanisms of stress-induced depression, particularly the lack of measurable biochemical and molecular alterations, hamper effective pharmacological interventions to prevent or ameliorate the behavioral deficits.

Therefore, the goal of this work is to we examine the distinctive behavioral impacts and neurobiological mechanisms of in utero exposure alone (prenatal one-hit stress) versus impaired maternal care alone (postnatal one-hit stress) or combined exposure (prenatal and postnatal two-hits stress). We hypothesized that prenatal stress acts as a priming factor that synergistically interacts with subsequent early-life challenges (inadequate caregiving by traumatized mothers) to exacerbate the tendency for offspring of traumatized mothers to develop psychiatric disorders later in life.

To recapitulate intergenerational trauma experience, we modeled stress by exposing mouse dams in the last four days of their pregnancy to a predator odor. We applied cross-fostering procedures to evaluate the distinct contributions to adult behavioral deficits of prenatal trauma and subsequent postnatal inadequate caregiving. We used a comprehensive “omics” approach combining metabolomic, transcriptomic, and bioinformatic analyses to determine the molecular processes that are associated with stress-induced behavioral deficits. Based on our findings of the mechanisms involved, which revealed a role for mitochondria metabolism dysfunctions, we examined whether the behavioral deficits can be reversed by early and/or late supplementation with acetyl-L-carnitine (ALCAR).

## Methods and Experimental Designs

### Animals and Breeding Procedures

Eight-week-old Swiss Webster mice were obtained from Charles River Laboratories (Wilmington, MA). One male and two female mice were group-housed, and pregnant mice were individually housed from the fourteenth day of pregnancy (gestational day 14) until delivery. After weaning (postnatal day 21), male mice were group-housed separately (Fig. 1a). All experimental procedures were approved by the Institutional Animal Care and Use Committee of University of California, Irvine and were performed in compliance with national and institutional guidelines for the care and use of laboratory animals.

**Figure 1.**
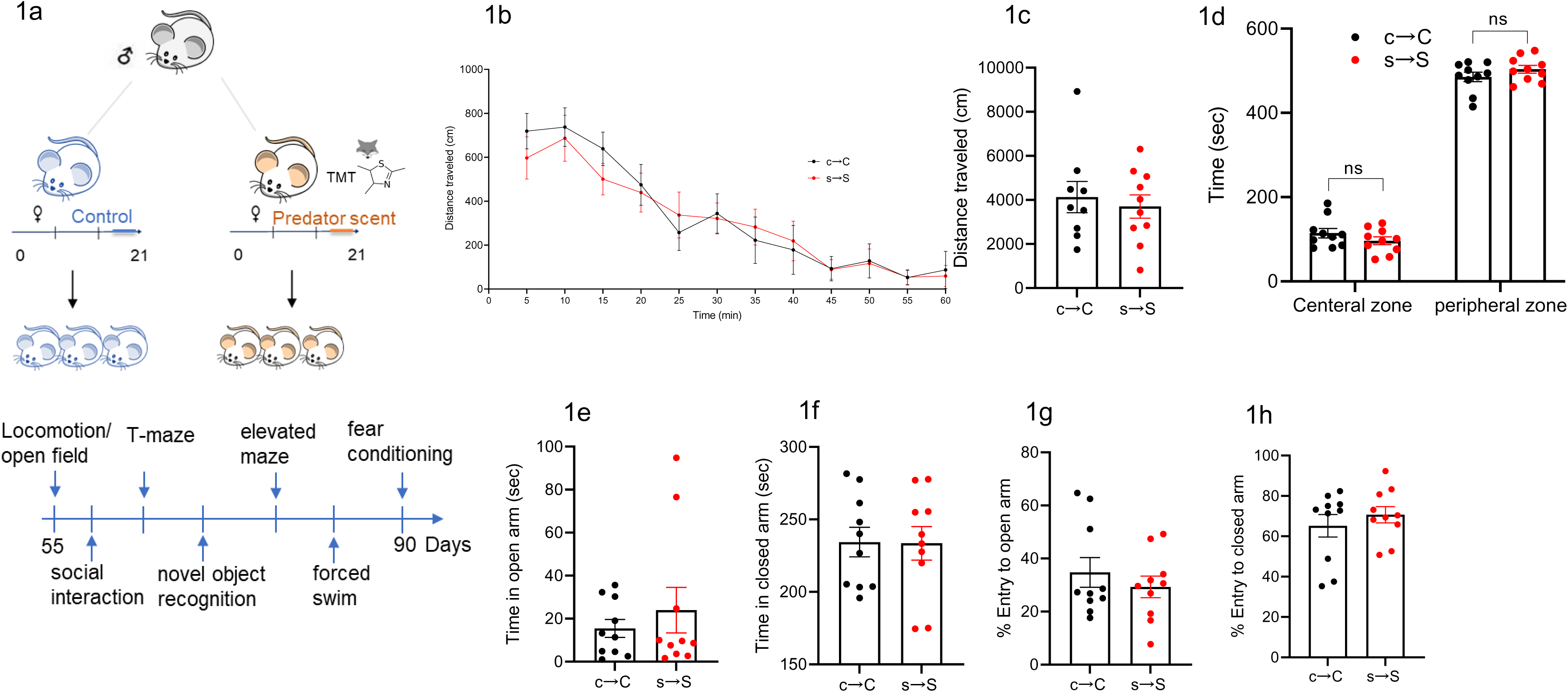
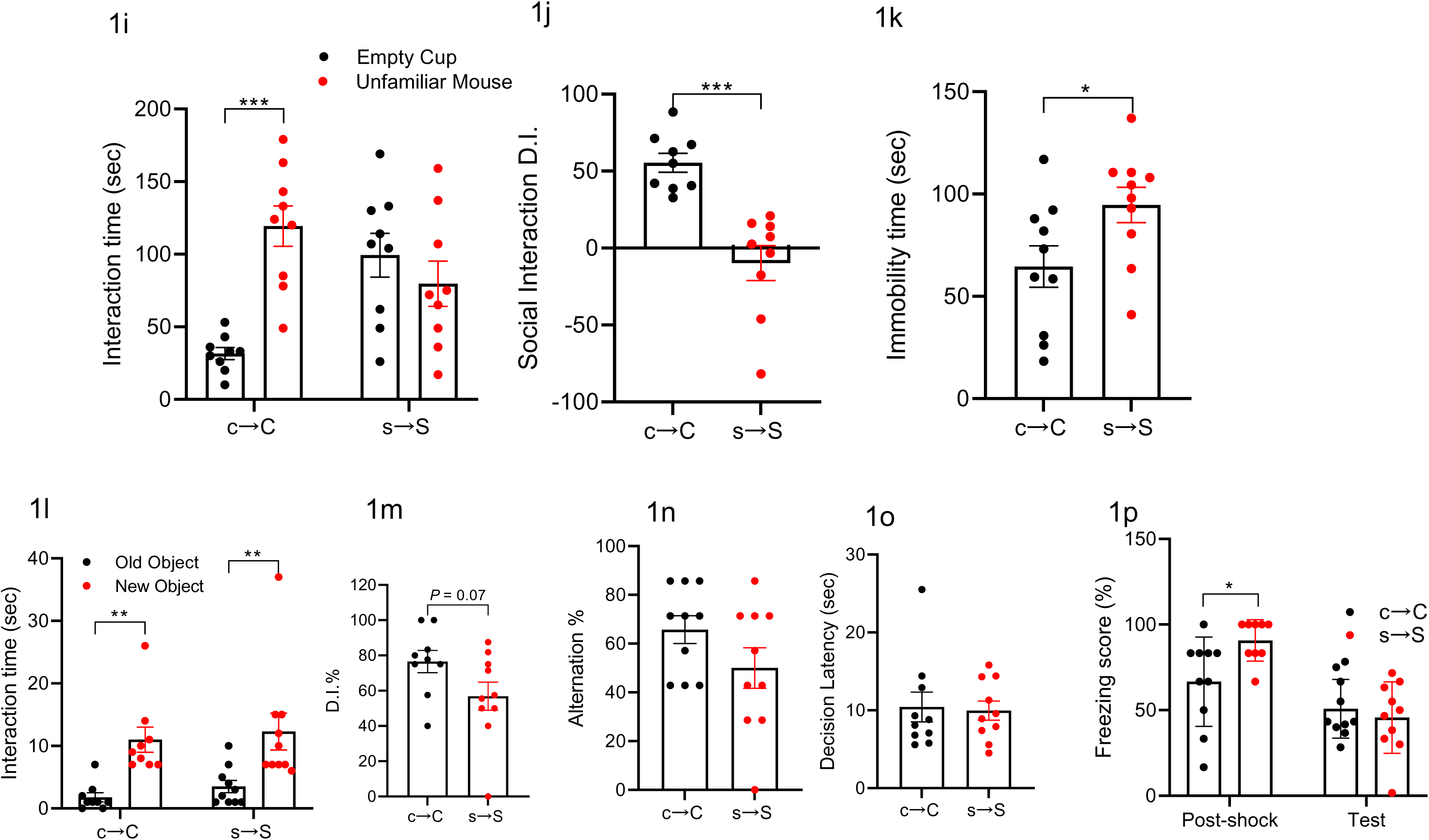
The s→S pups display depressive-like behavior, social impairment, but normal cognitive functions. **a.** Schematic of the experiment design **b,c.** s→S pups display normal locomotor activity. **(b)** Time-course of the locomotor activity, **(c)** Distance mice travelled in 60 minutes of the locomotion assay (n=10 c→C, 10 s→S). Unpaired student test (t=0.2723, *P=*0.7885): c→C vs s→S, ns, not significant. **d.** Time mice spent in the central and peripheral zones in 10 minutes in the locomotion assay (n=10 c→C, 10 s→S). Two way ANOVA revealed a significant zone effect (F_1,36_=1453, *P<*0.0001) followed by Bonferroni post-hoc test: c→C vs s→S, ns, not significant. **e,f.** Elevated plus maze measure of percent entry to **e.** closed (unpaired t-test; t=0.788, *P=*0.44), and **f.** open arms (unpaired t-test; t=0.788, *P=*0.44). **g,h**. Elevated plus maze measure of time spent in **g.** closed (unpaired t-test; t=0.05, *P=*0.95), and **h.** open arms (unpaired t-test; t=0.74, *P=*0.46). **i.** Time mice spent interacting with empty cup and unfamiliar mouse in the social interaction assay (n=9 c→C, 9 s→S). Two way ANOVA: stress effect (F _1,32_=6.783, *P=*0.0138), stress x object interaction (F_1,32_=16.88, *P=*0.0002) followed by Bonferroni post-hoc test: empty cup vs unfamiliar mouse, **** *P<*0.0001, ns, not significant. **j.** Discrimination index in the social interaction assay (n=9 c→C, 9 s→S). Unpaired student test (t=5.053, *P=*0.0001): c→C vs s→S, *** *P<*0.001. **k.** Time mice spent immobile in the forced swim assay (n=10 c→C, 10 s→S). Unpaired student test (t=2.27, *P=*0.0358): c→C vs s→S, * *P<*0.05. **l.** Time mice spent exploring both the new and old objects during the test session in the novel object recognition assay (n=9 c→C, 10 s→S). Two-way ANOVA revealed a significant stress effect (F _1,34_=21.79, *P<*0.0001) followed by Bonferroni post hoc test: old object vs new object, ** *P<*0.01. **m.** Discrimination index in the novel object recognition assay (n=9 c→C, 10 s→S). Unpaired student test (t=1.904, *P=*0.074): c→C vs s→S, ns, not significant. **n.** Percentage of the alternation choice mice made in the T-maze spontaneous assay (n=10 c→C, 10 s→S). Unpaired student test (t=1.558, *P=*0.1366): c→C vs s→S, ns, not significant. **o.** Time mice spent to make a decision in T-maze spontaneous assay (n=10 c→C, 10 s→S). Unpaired student test (t=0.2029, *P=*0.8415). **p.** Percentage of freezing behavior in contextual fear conditioning assay (n=10 c→C, 10 s→S). Two way ANOVA revealed a significant stage effect (F_1,35_=22.82, *P<*0.001) and stress x stage interaction (F_1,35_=5.260, *P=*0.0279) followed by Bonferroni post hoc test: c→C vs s→S, \**P<*0.05, ns, not significant.

### Predator Scent (PS) Exposure and Cross-fostering

On gestational day 17, the pregnant mice were exposed to a predator scent (2,5-dihydro-2,4,5- trimethylthiazoline, or TMT, a constituent of fox urine) for 1 hour on each of four consecutive days in a separate cage under the fume hood and were then returned to their home cage. Bedding mixed with a few drops of distilled water (Control, C) or 0.02% TMT (Stress, S) was placed in a 60 x 15 mm polystyrene plate placed in each cage to achieve scent exposure [14–17].

### Cross-fostering mouse model and animal care

Pups (control, c or stress, s) of control (C) and stress (S) mothers were raised by their biological mother (c→C and s→S) or were cross-fostered to the opposite mother treatment (s→C and c→S, for control mothers with stress pups or stress mothers with control pups, respectively) within 24- hr of birth. The pups remained with either their biological or foster mother until the weaning day (postnatal day 21). Subsequently, the male mice were then selected for the study and group-housed in groups ranging from 3 to 5.

### Behavioral analyses

Male mice were tested from postnatal week 8 to week 13 with a battery of behavioral paradigms in the following order: locomotion and stereotypy/open field, social interaction, spontaneous T-maze alternation, novel object recognition, elevated plus maze, forced swim, contextual fear conditioning (Fig. 1a). The sequence of specific assays spaced by 3-6 days inter-assay interval was adapted from previous reports [18, 19]. In subsequent cross-fostering behavioral studies, mice were tested only in social interaction and forced swim assays. In ALCAR behavioral studies, mice were tested only in one behavioral assay (forced swim).

#### Locomotor Activity

Mice were placed into a locomotion test chamber (Med Associates, Inc.) for 90 minutes as we previously described [20]. The first 30 minutes was allotted for the animal to habituate to the chamber before the test. The horizontal, vertical, and stereotypic activities were then recorded for the remaining 60 minutes and were then analyzed by the Activity Monitor 5 software (Med Associates, Inc.).

#### Open Field Assay

The open field assay was carried out as described previously with slight modifications [21]. Prior to testing, the open field test chamber (40 x 40 cm, Med Associates, Inc.) was sectioned into a central zone, a 24 x 24 cm square in the middle of the test chamber, and a peripheral zone, the remaining outer area. Mice were placed in the chamber and the time the animal spent within 10 minutes in the central or peripheral zones was recorded. Using the predetermined zone areas, the center-to-periphery exploration time ratio was assessed by the Activity Monitor 5 software (Med Associates, Inc.).

#### Elevated Plus Maze

A standard elevated plus-maze, made of grey Plexiglas, was placed in a sound-proof observation room with controlled light (200 Lux) on the central platform of the maze [22]. Animals were given a 30-minute period to habituate to the room before being tested. During the test, the mice were placed in the center of the plus facing an open arm and were given 5 minutes to explore. The behavior was recorded and scored by two independent observers blind to the animal treatments. The animals were scored based on time spent in the closed and open arms and the number of entries to the closed and open arms.

#### Social Interaction

The social interaction test was performed using the three-chambered apparatus as previously described [21]. Two wire mesh cups were placed in the middle of both the right and left chamber (one on each side). In one of the cups on either side, a control mouse (an unfamiliar mouse of the same strain, gender, and age with no prior contact with the subject mouse) was placed. The subject mouse was placed in the middle chamber and was given 5 minutes to explore with the dividing doors closed. The dividing doors were then removed to allow the mouse to freely travel between the 3 chambers for 10 minutes. The duration of direct contact between the subject mouse and both the empty cup and the cup with the mouse was measured. The relative exploration time was recorded and expressed by a discrimination index: [D.I. = (*T_mouse_ – T_empty_)/ (T_mouse_ + T_empty_*) x 100%]. The test was video recorded and analyzed by the ANY-MAZE software (Stoelting Co.).

#### Spontaneous T-Maze Alternation Assay

The spontaneous T-maze alternation assay was carried out as described previously [21]. The T-maze has three designated parts: the main stem, the side arms, and the starting area (AccuScan Instruments, Inc.). The mice were placed in the starting area behind a sliding door, blocking the main stem and side arms and were given a 30 second acclimation period before the start of each trial. After the acclimation period, the sliding door was removed, and the mice can choose either the left or right-side arm. After a choice (all four paws in the chosen arm), the sliding door was closed behind the animal, allowing the mouse to explore the chosen arm for 30 seconds, before being manually returned to the start area for the next trial. A total of eight trials were completed, allowing for a total of seven total possible alternations. The alternation percentage was calculated as 100 x (number of alternations/7). The time for the animal to decide on a side arm (decision latency) was also recorded.

#### Novel Object Recognition Assay

The novel object recognition (NOR) assay was performed as described previously [20]. The assay consists of training and testing phases. Prior to training, the mice were handled for 1-2 minutes a day for 3 days and were given 10 minutes a day for 3 days inside the empty experimental apparatus to habituate to the environment. During the training phase, mice were placed in the experimental apparatus, and were given 10 minutes to explore and examine two identical objects. Twenty-four hours later, one of the familiar objects was replaced with a novel object and the animals were given 5 minutes to explore the chamber. During both the training and testing phases, the duration and number of times the mice interact with the familiar and novel objects were recorded individually. The relative exploration time was recorded and expressed by a D.I.: [D.I. = (*T_novel_ - T_familiar_)/ (T_novel_ + T_familiar_*) x 100%]. Tests were video recorded and analyzed by the ANY-MAZE software (Stoelting Co.).

#### Forced swim

The forced swim assay was performed as previously described [23]. During the 6-minute assay, the first 2 minutes of activity was discarded. For the last 4 minutes, mice were videotaped and the immobility time was recorded. The ANY-MAZE software was used to record and analyze immobility (Stoelting Co.).

#### Contextual Fear Conditioning Assay

The Contextual Fear Conditioning Assay was performed as previously described [20]. The assay was split into a training and retention session. Prior to the training session, mice were handled for 1-2 minutes per day for 3 days. During the training session, mice were placed in the conditioning chamber (TSE Systems, Inc.), with a striped pattern, for 2.5 minutes, before receiving a 0.7 mA foot shock for 2 seconds. After an additional 30 seconds, the mice were returned to their home cages. The freezing behavior (lack of movement for at least 3 seconds in a 5 second interval) pre-shock and post-shock were measured. Twenty-four hours after training, the mice were placed in the chamber again, with the same striped pattern, for 5 minutes and the freezing behavior, without shock, and was assessed. Freezing behavior was scored as freezing (1) or not (0) for every 5 second interval and the percentage of freezing behavior was calculated as 100 x (number of freezing intervals/total intervals).

### Acetyl-L-carnitine (ALCAR) Treatment

The ALCAR treatment protocol was adapted from a previously reported protocol [24]. The s→S mice received ALCAR (0.3%, Sigma Aldrich) in the drinking water either on postnatal day 21 (PPD 21 weaning day) and continued to week 8, or on week 7 and continued to week 8. Control groups received normal tap water. Forced swim tests were carried out on the last day of the treatment and 7 days after discontinuing ALCAR.

### Immunohistochemistry

Immunohistochemistry methods to examine the Arc positive (Arc+) neurons were conducted as we previously described [23]. Mice were perfused under isoflurane anesthesia with 4% paraformaldehyde (PFA), and brains were removed, post-fixed in 4% PFA for overnight at 4°C, and stored in 30% sucrose. Coronal sections were cut at 20 µm thickness, and three sections were used from each region of interest. Brain regions were defined according to their anatomy using the mouse brain atlas [25]. Sections were blocked with 4% normal goat serum in PBS with 0.3% Triton X-100 for 1 hour, and were then incubated in the blocking buffer that contains the primary antibodies (Arc 1:500, Sigma). The sections were then washed with PBS, incubated with the secondary antibody (1:500) for 1 hour, then with 4’,6-diamidino-2-phenylindole (DAPI) for 5 minutes, and were then mounted on slides. Imaging was carried out using Leica Sp8 TCS confocal microscope (UCI optical biology core facility). Arc positive (Arc^+^) neurons were counted in the bilateral areas of each section, and the mean counts of three non-consecutive sections per brain of 3-5 brains were calculated. Cell counts were performed using ImageJ [26], and confirmed manually by two persons blind to the experiment conditions.

### Brain metabolite analyses

Tissues of one brain hemisphere (cortical-subcortical/mouse) were collected from neonatal and adult mice, and extracted following the protocols first published in [27]. Metabolic profiling was carried out by the West Coast Metabolomics Center (University of California Davis). Three metabolic platforms were profiled: 1) primary metabolites including hydroxyl acids, purines, pyrimidines, carbohydrates and sugar phosphates, amino acids, and aromatics; 2) lipids; and 3) biogenic amines and methylated and acetylated amines [28].

### mRNA Microarray Analysis

Microarray experiments and analysis were performed as we previously described [20, 29]. Total RNA was extracted (Qiagen RNA extraction kit) from one brain hemisphere of the 24-hours’ and 13 weeks’ old pups according to the manufacturer’s protocol. RNA samples with A260/A280 absorbance ratios between 2.00-2.20 were reverse-transcribed into cDNA and analyzed by “whole-transcript transcriptomics” using the GeneAtlas microarray system (Affymetrix) and manufacturer’s protocols. MoGene 2.1 ST array strips (Affymetrix) were used to hybridize to newly synthesized ss-cDNA. Each array comprised 770,317 distinct 25-mer probes to probe an estimated 28,853 transcripts, with a median 27 probes per gene. Only annotated genes were included in the differential analysis.

### Real time quantitative polymerase chain reaction (RT-qPCR)

RNA was reverse-transcribed into cDNA using the Transcriptor First Strand cDNA synthesis kit (Roche Molecular Systems, Inc) following manufacturer’s instructions. RT-qPCR was performed using the LightCycler480 SYBR Green I Master kit (Roche Molecular Systems, Inc), and analyzed using the amplification reactions were carried out on a LightCycler® 480 instrument with gene specific primers listed in the supplementary methods. Samples from 5 animals were run in duplicates and averaged. The comparative CT method was used to obtain the Ct values from each RNA sample and normalized to a housekeeping gene (GAPDH) [30]. The difference between the Ct values (ΔCt) of the gene of interest and the housekeeping gene was calculated for each sample and the difference in the ΔCt values between the experimental and control samples ΔΔCt was calculated. Fold change was obtained as 2^(-ΔΔCt) [20].

### Bioinformatic analysis of metabolomics and transcriptomics, and metabolites-transcriptomic Integration

The bioinformatic analysis of the differential metabolites differential genes was conducted as we described previously [28]. A differential analysis was performed between the s→S and c→C groups for both 24-hour old pups and 13-week old adult mice groups using the Cyber-T program [31, 32] to identify the top up- or down-regulated metabolites, using p-value cutoff at 0.05. The metabolic pathway enrichment analysis was performed on the differentially expressed metabolites using Fisher exact test, which is based on a hypergeometric distribution, to identify enriched pathways from the Small Molecule Pathway Database [119, 120]. The inputs of the 2×2 contingency table used for the Fisher exact test were: number of metabolite hits, number of metabolite non-hits, number of non-hits in metabolites associated with the pathway, and number of background metabolites. Pathways with p-value less than 0.05 were considered enriched in the given list of metabolites. Differentially expressed genes in s→S and c→C groups for both neonatal and adult mice were identified from the transcriptomic data and underwent further analyses. These analyses were: (1) pathway enrichment analysis using fisher exact test on pathways from the Pathway Common Database [33] and the ConsensusPathDB [34], and STRING analysis (https://string-db.org/); (2) upstream transcription factor binding site (TFBS) enrichment analysis of promoters using the promoters and their target genes identified by the MotifMap database [35, 36] and the CHiPSeq database from ENCODE [37]; (3) identification of transcription factors (TFs) and RNA-binding proteins (RBP) and analyses of their differentially expressed downstream targets using MotifMap and CHiPSeq databases.

For Integrated metabolite-transcriptomic analysis, metabolites were identified by InChI Key or name in Human Metabolome Database (HMDB) [38]. Enzymes and enzyme-coding genes involved in metabolic reactions of the metabolites were queried from the biological properties section of the database. The transcriptomic expression of those enzyme-coding genes in s→S and c→C groups was extracted from the microarray data. In this way, the activity of the metabolites and their enzyme-coding genes in the s→S and c→C groups were linked together for holistic analysis. Network graphs were constructed with nodes being a metabolite or a gene, and edges being a known reaction between the metabolite and enzyme coded by the gene.

### Integrated mouse-human transcriptomic analysis

The DEGs in both adult and newborn mouse groups from the transcriptomic analysis were compared with DEGs in human depression [39]. The overlaps between DEGs common to stress mice and human depression were computed for adult and newborn mouse groups using Fisher exact test.

### Statistical analysis

Statistical analyses of behavioral data were carried out using GraphPad Prism (GraphPad Software, Inc.). Data were presented as means ± S.E.M. Results were analyzed by unpaired student t-test, one-way, and two-way ANOVA followed by the appropriate post hoc comparisons, and *P<*0.05 was considered statistically significant.

## Results

### s→S pups display depressive-like behavior, social impairment, but normal cognitive functions

Compared to the offspring of control dams (c→C), the offspring of dams exposed to the predator scent (s→S) exhibited normal locomotor activities (Fig. 1b,c). The c→C and s→S mice travelled similar distances in the center and peripheral area in the open field assay (*P*>0.05, unpaired t-test, Fig. 1d), and displayed similar times and number of entries to the open arm of the elevated maze (*P>*0.05, unpaired t-test, Fig. 1e-h), indicating normal anxiety behavior in the s→S mice. In the social interaction assay, s→S mice displayed less interaction with the unfamiliar mice than did the c→C group (*P<*0.001, two way ANOVA, followed by Bonferroni post-test, Fig. 1i), and their D.I. was significantly lower (*P<*0.001, unpaired t-test, Fig. 1j), indicating impaired sociability in the s→S mice. The s→S mice displayed depressive-like behavior, reflected by their higher immobility time compared to the c→C mice (*P<*0.05 unpaired t-test, Fig. 1k).

We investigated whether the prenatal stress caused cognitive dysfunction, by testing object memory, working memory, and contextual fear memory. In the novel object recognition assay, both groups spent more time exploring the new object than the old object (*P<*0.01, two-way ANOVA, followed by Bonferroni post-test, Fig. 1l), the D.I. in the s→S mice was not different compared to the c→C mice (*P*=0.07, unpaired t-test, Fig. 1m). In the spontaneous T-maze alternation assay, s→S and c→C mice showed a similar percentage of arm choice alternation and decision latency (*P>*0.05, unpaired t-test, Fig. 1n,o). In the contextual fear-conditioning assay, the s→S group exhibited higher freezing behavior during the stimulus session (*P<*0.05), but not in the retention session (*P>*0.05, two-way ANOVA, followed by Bonferroni post-test, Fig. 1p). Together, these results suggest that s→S mice display normal cognitive function. Mothers exposed to predator scent exhibited deficits in maternal behavior, and depressive-like behavior, but not locomotor activity or anxiety behavior (Fig. S1a-g).

### Prenatal stress results in alterations in brain metabolomics

Global brain metabolomics analysis of neonatal pups revealed that prenatal exposure to stress induced alterations in 50 brain molecules (Fig. 2a), of which the mitochondrial metabolite 2- hydroxyglutaric acid (2-HG) displayed the highest increase in the brains of neonatal s→S pups (Fig. 2b, Table S1). 2-HG is a hypoxia and mitochondrial dysfunction marker, and an epigenetic modifier [40–43]. Therefore, its striking increase in the brains of the s→S pups within the first 24 hours after birth indicates disruptions in mitochondrial respiratory functions and epigenetic processes. Alongside 2-HG elevated levels, the levels of tricarboxylic acid (TCA) cycle metabolites succinic acid and γ-aminobutyric acid (GABA) significantly increased in the brains of the neonatal s→S pups (Fig. 2b,c, Table S1). Beside its role as a neurotransmitter, GABA regulates the TCA cycle through the GABA shunt [44], in which 2-HG is formed as a by-product during conversion of GABA to succinic acid (Fig. 2c) [45–49]. The increases in 2-HG, GABA, and succinate levels further support a mitochondrial metabolism dysfunction, particularly in the TCA cycle and GABA shunt, in the neonatal s→S pups.

**Figure 2.**
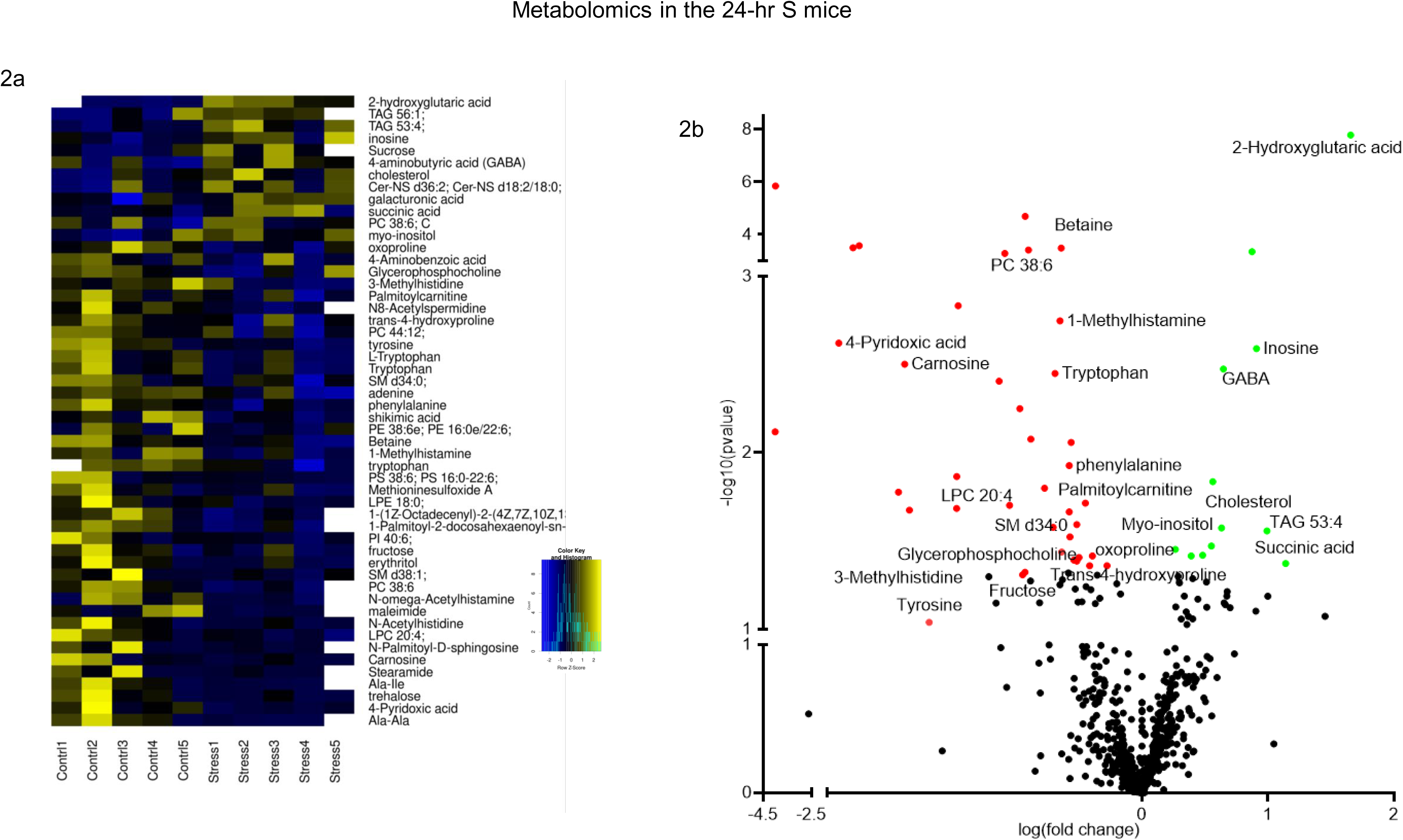
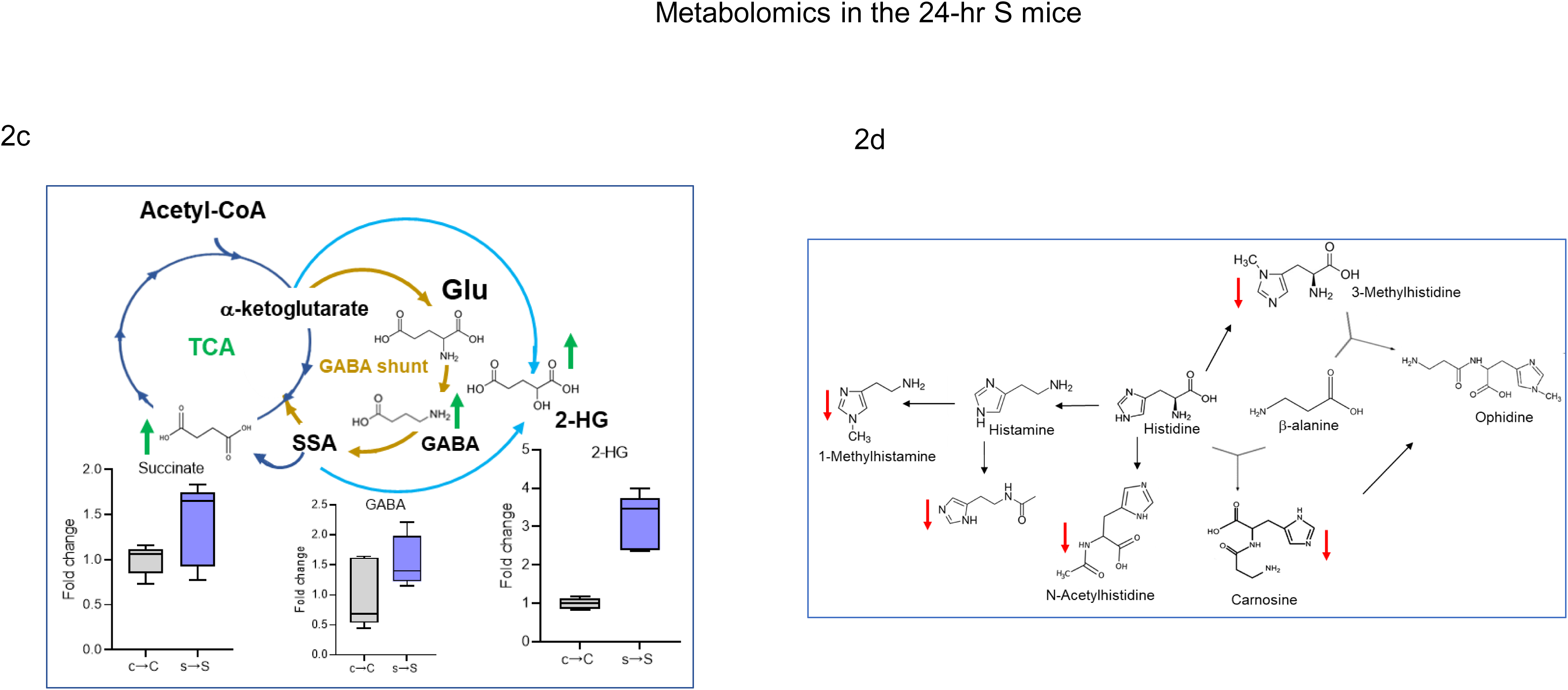
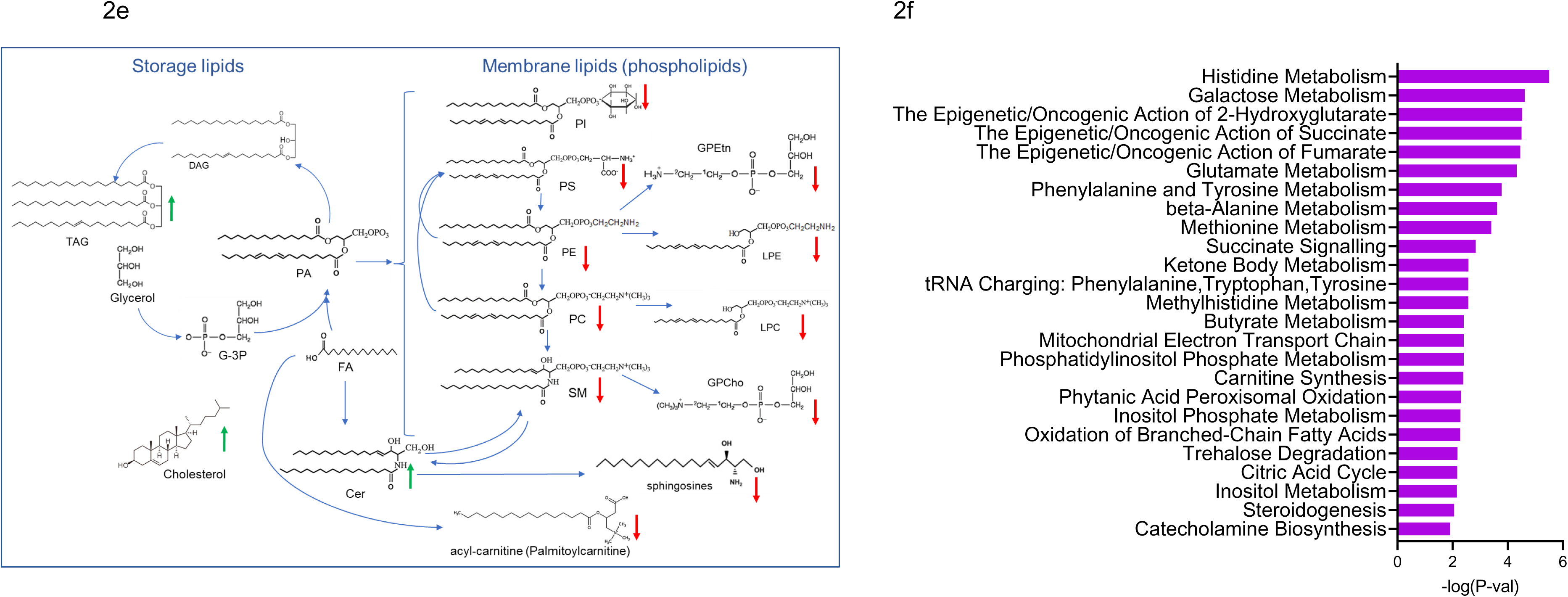
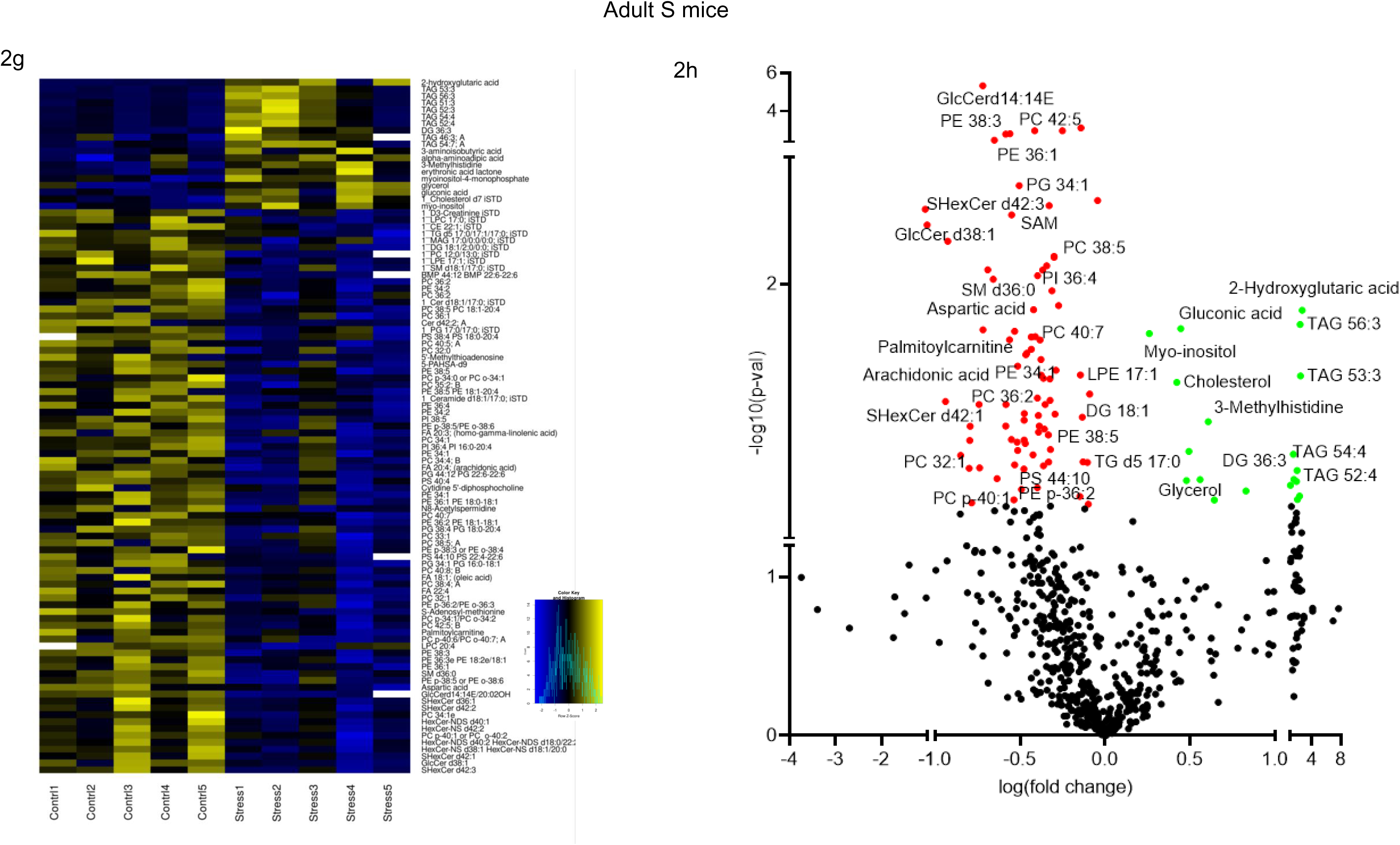
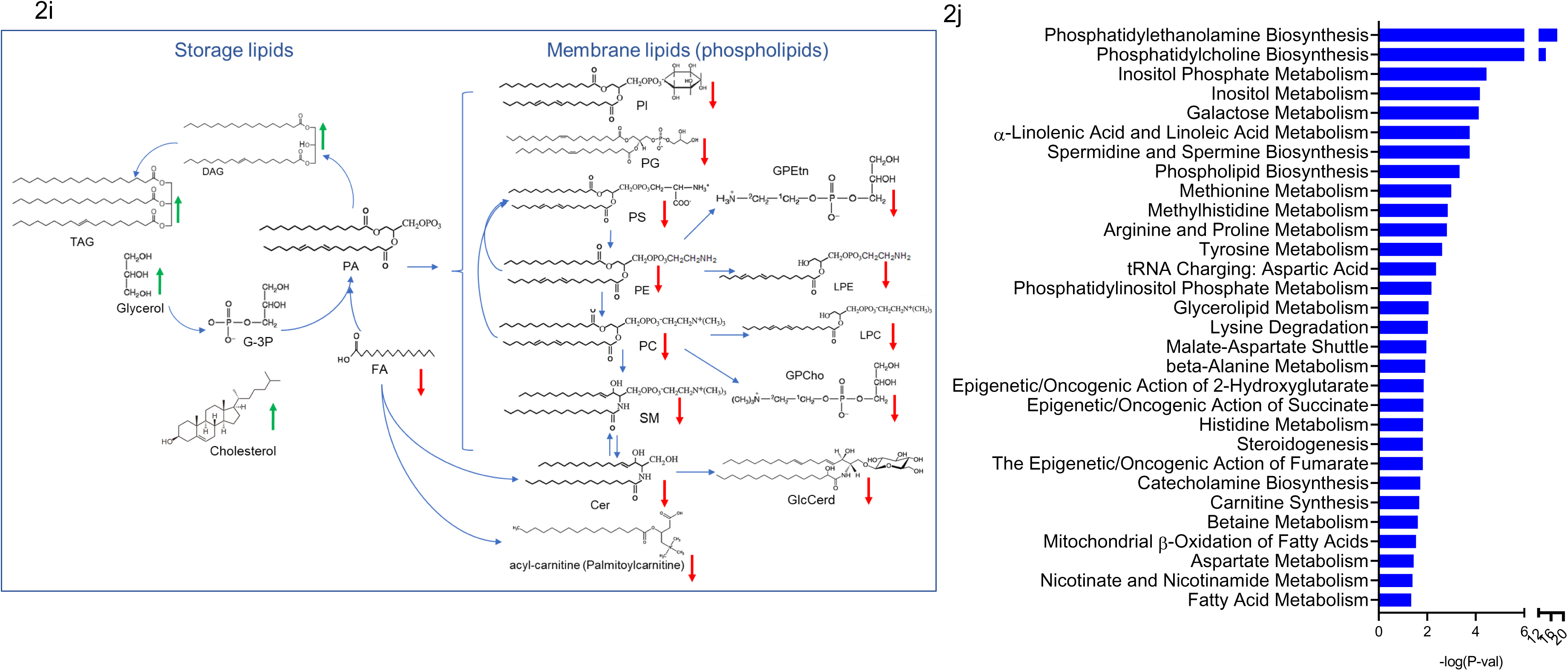
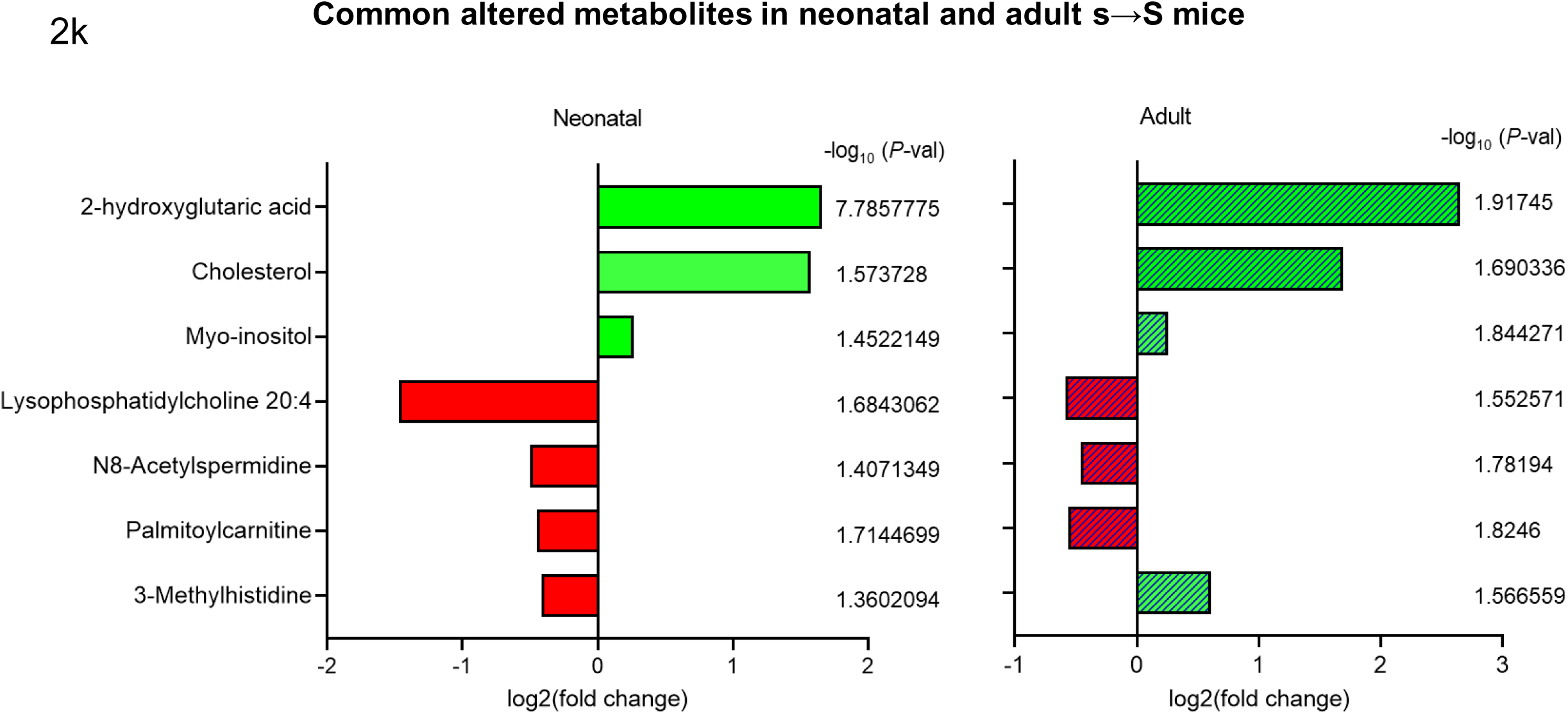
Prenatal exposure to stress causes changes in brain metabolites. **a.** Heatmap of metabolites significantly changed in the brains of neonatal s→S mice (*P*<0.05). **b.** Volcano plot of metabolites significantly changed in the brains of neonatal s→S mice (*P*<0.05). **c.** Effect of prenatal exposure to stress on the tricarboxylic acid (TCA) and GABA shunt metabolism. Increased metabolites in green. **d.** Changes of metabolites related to histidine metabolism, decreased metabolites in red. **e.** Changes of metabolites related to lipid synthesis/metabolism in the neonatal s→S pups. Increased metabolites in green, and decreased metabolites in red. Phosphatidylcholines (PC), lysophosphatidylcholines (LPCs), phatidylethanolamines (PE), lysophosphatidylethanolamines (LPE), phosphatidylserine (PS), fatty acids (FA), arachidonic acid, glucoceramides (GlcCer), Phosphtatidylglycerols (PG), and the decrease in sphingomyelins (SMs), diacylglycerols (DAG), and triglycerides (TAGs) **f.** Pathway analysis of metabolites significantly changed in the neonatal MET pups (Table S2 shows details of number of compounds, hits and p values) **g.** Heatmap of metabolites significantly changed in the brains of adult s→S mice (*P*<0.05). **h.** Volcano plot showing differentially expressed metabolites in the brains from adult s→S mice. **i.** Changes of metabolites related to lipid synthesis/metabolism in adult s→S pups. Increased metabolites in green, and decreased metabolites in red. **j.** Pathway of metabolites significantly changed in the adult s→S pups (Table S2 shows details of number of compounds, hits and p values). **k,** Metabolites differentially expressed in neonatal and adult s→S mice with log_2_(fold change) and –log10 (P-value)

Other metabolites that were altered in the brains of the neonatal s→S pups include molecules involved in glycolysis and energy metabolism such as *myo*-inositol (increased) and palmitoyl-carnitine (decreased) (Fig. 2b Table S1). The decrease in palmitoyl-carnitine reflects reduced oxidative capacity and mitochondrial ATP production [50–53]. The mitochondrial dysfunction in neonatal s→S pups is further indicated by the marked decrease of carnosine (> threefold) [54, 55], which was associated with decreases of other histidine metabolism components (Fig. 2b,d, Table S1,2). Carnosine (β-alanyl-L-histidine) is a dipeptide that has mitochondrial pH-buffering capacity and enhancing activity on the respiratory chain complexes, and mitochondrial energy production [54–57]. The depletion of these energy-related molecules, thus, suggests that the oxygen-glucose deprivation in the neonatal s→S pups’ brains creates a condition of excessive need for carnosine and its derivatives, to restore glycolysis and ATP production, and that the need exceeds the resources available of these molecules [58]. On the other hand, carnosine depletion might result in a reduction in the buffering of mitochondrial pH, thus contributing to the cellular and particularly mitochondrial acidity, associated with the production of 2-HG [41].

Metabolites of monoamine neurotransmitters’ pathways displayed decreases in the brains of neonatal s→S pups including noradrenaline and dopamine precursor (tyrosine), serotonin precursor (tryptophan), and histamine-metabolism components (Fig. 2b).

The major lipidomic changes included increases in triacylglycerols (TAGs), cholesterol, ceramides (Cer), and decreases in acyl-carnitine (palmitoyl-carnitine), and membrane phospholipids (Fig. 2a,e). Bioinformatic analysis of metabolites altered in the brains of neonatal s→S pups substantiated enrichment of pathways associated with epigenetic processes, energy metabolism, mitochondrial functions, fatty acid oxidation, and complex lipid metabolism (FDR < 0.05, Fig. 2f, Table S1).

In the adult brains, 2-HG again exhibited the highest change (> 6-fold increase) in the brains of the s→S pups, indicating mitochondrial dysfunction (Fig. 2g-h, Table S1). Other metabolites whose levels increased are molecules involved in energy and lipid metabolism/storage (myo-inositol, glycerol, TAGs, and cholesterol). There were profound abnormalities in the membrane, mitochondria, and signaling lipidomics. Except for TAGs and diacylglycerol (DGs), which were increased in the S adult brains, all other lipids were decreased (Fig. 2h-i). Among the notable metabolites whose levels decreased in the s→S mice were aspartic acid, palmitoyl-carnitine, and S-Adenosyl-methionine (SAM).

Bioinformatic analysis revealed similar biochemical pathways enrichments in the brain’ metabolites of adult s→S mice to those significantly enriched in the neonatal s→S pups, indicating early but long-lasting metabolomic signatures produced by prenatal stress (FDR < 0.05, Fig. 2j, Table S2). This is further supported by the finding that seven metabolites exhibited changes in the same directions in both neonatal and adult s→S mice: 2-HG, *myo*-inositol, N8- Acetylspermidine, palmitoyl-carnitine, cholesterol, LPC 20:4, and methylhistidine (Fig. 2k). Hence, metabolic pathways enriched in the brains’ metabolites of both neonatal and adult s→S mice included epigenetic/oncogenic action of 2-HG and succinate, mitochondrial GABA metabolism, carnitine synthesis, mitochondrial beta-oxidation of long-chain fatty acids, catecholamine biosynthesis, glutathione metabolism, inositol metabolism, histidine metabolism, methionine metabolism, and phosphatidylinositol-phosphate metabolism (Fig. 2f,j).

### Prenatal stress results in distinct alterations in brain transcriptomics

Global brain transcriptomic analysis of differentially expressed genes (DEGs) revealed a subset of genes that exhibited a ≥ 1.5-fold change in the neonatal s→S mice (*P<*0.05); Avp, Egr1, C1ql1, Fos, Crhbp (downregulated) and Baiap2l1, mt-Tqm, Irak1bp1, and 11 microRNAs (Mirs) (upregulated) (Fig. 3a, Table S3). To verify the microarray results, we conducted RT-qPCR on six genes (Egr1, Fos, Avp, Vgf, Mcm6, and C1ql1) that were shown by microarray to display significant changes in the neonatal s→S mice (*P<*0.05); the RT-qPCR results confirmed the microarray findings (Fig. 3b).

**Figure 3.**
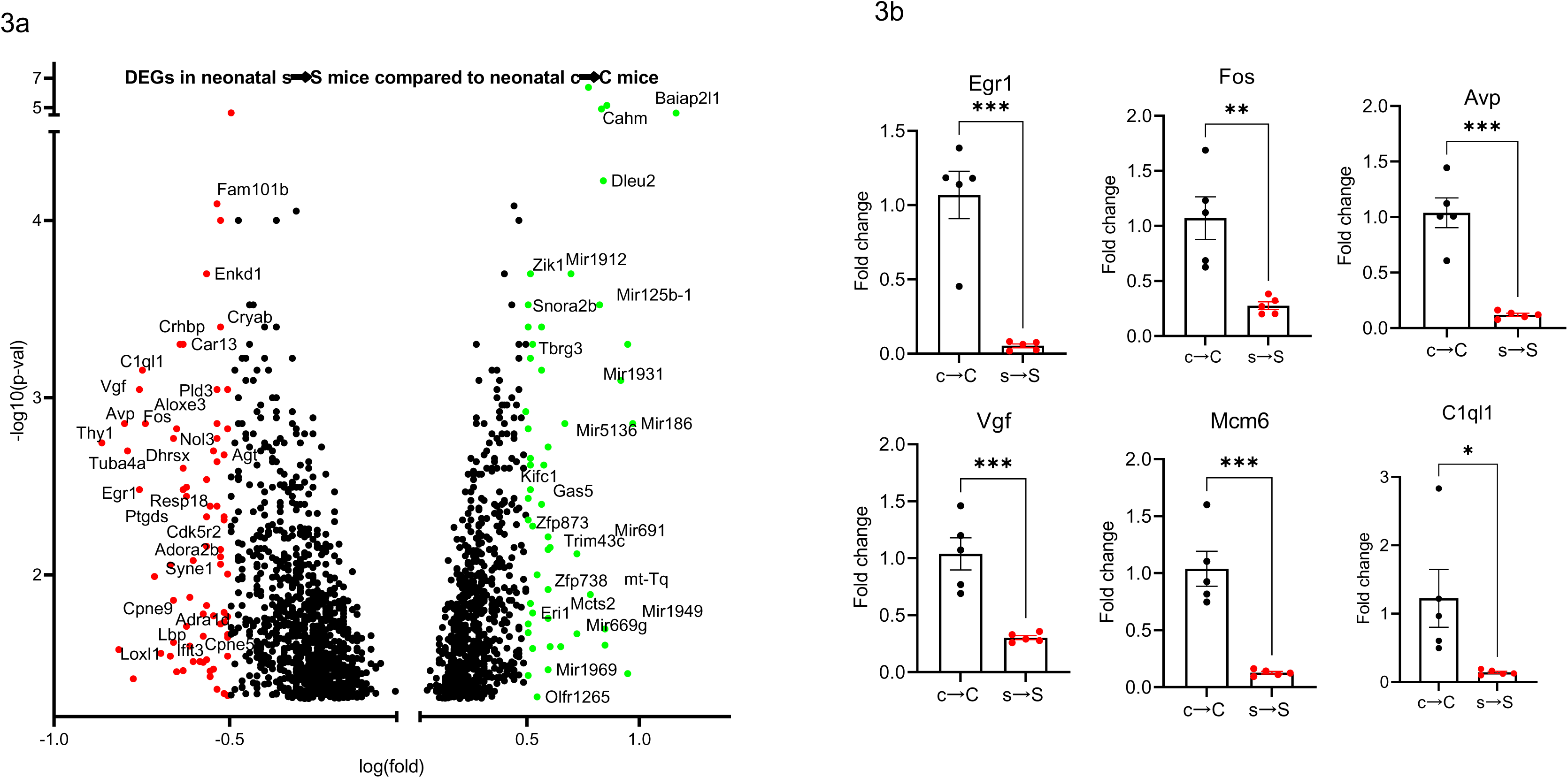
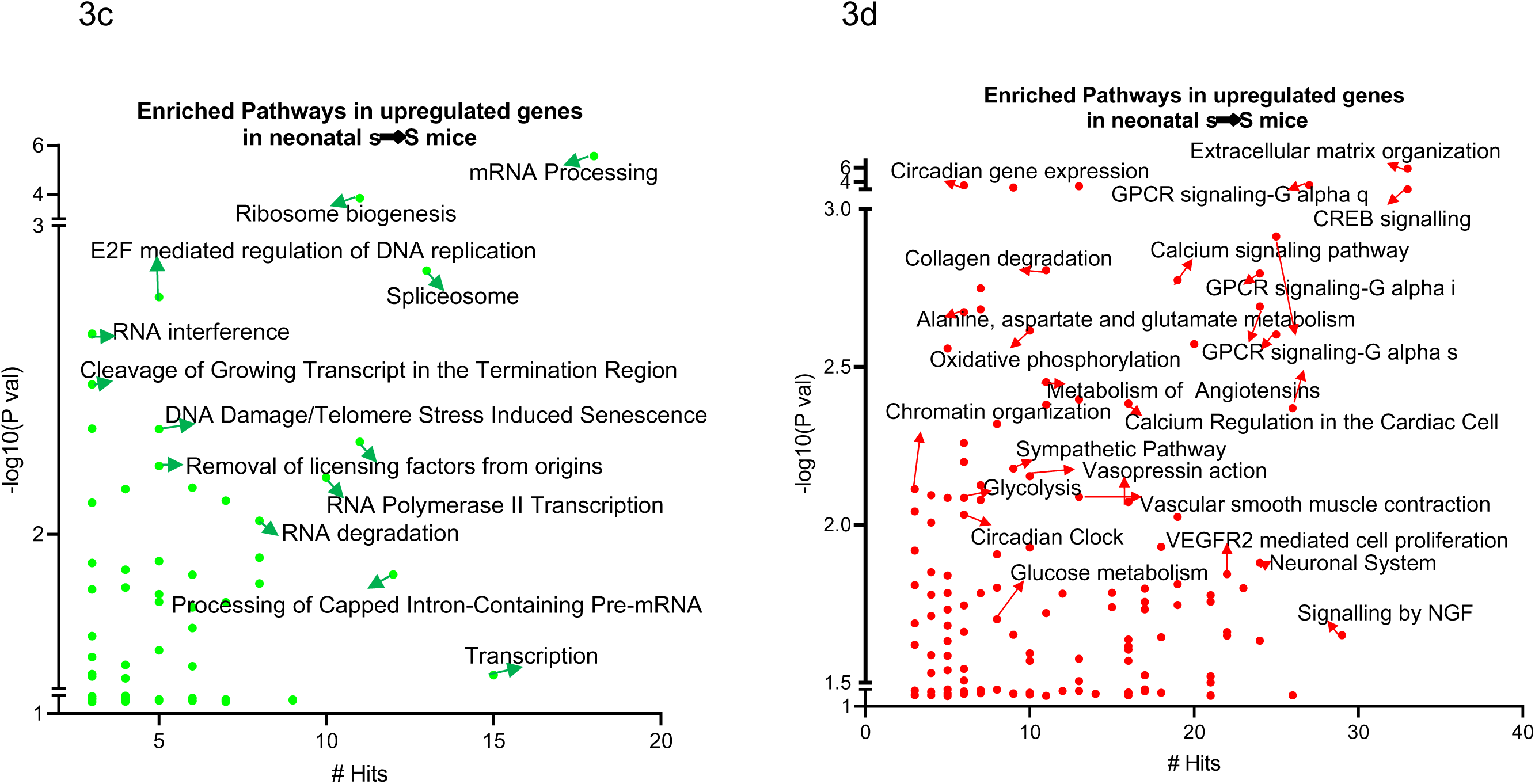
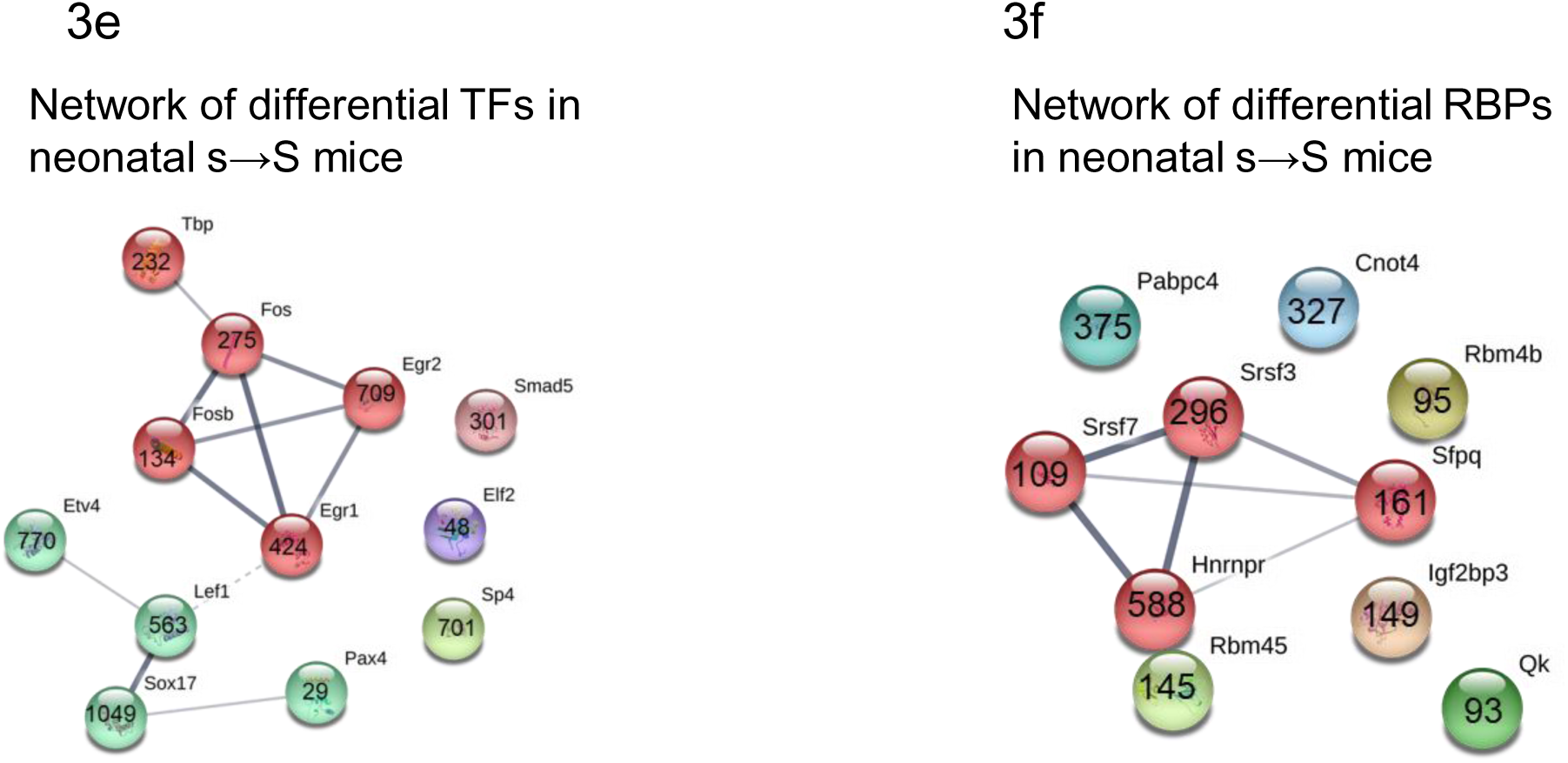
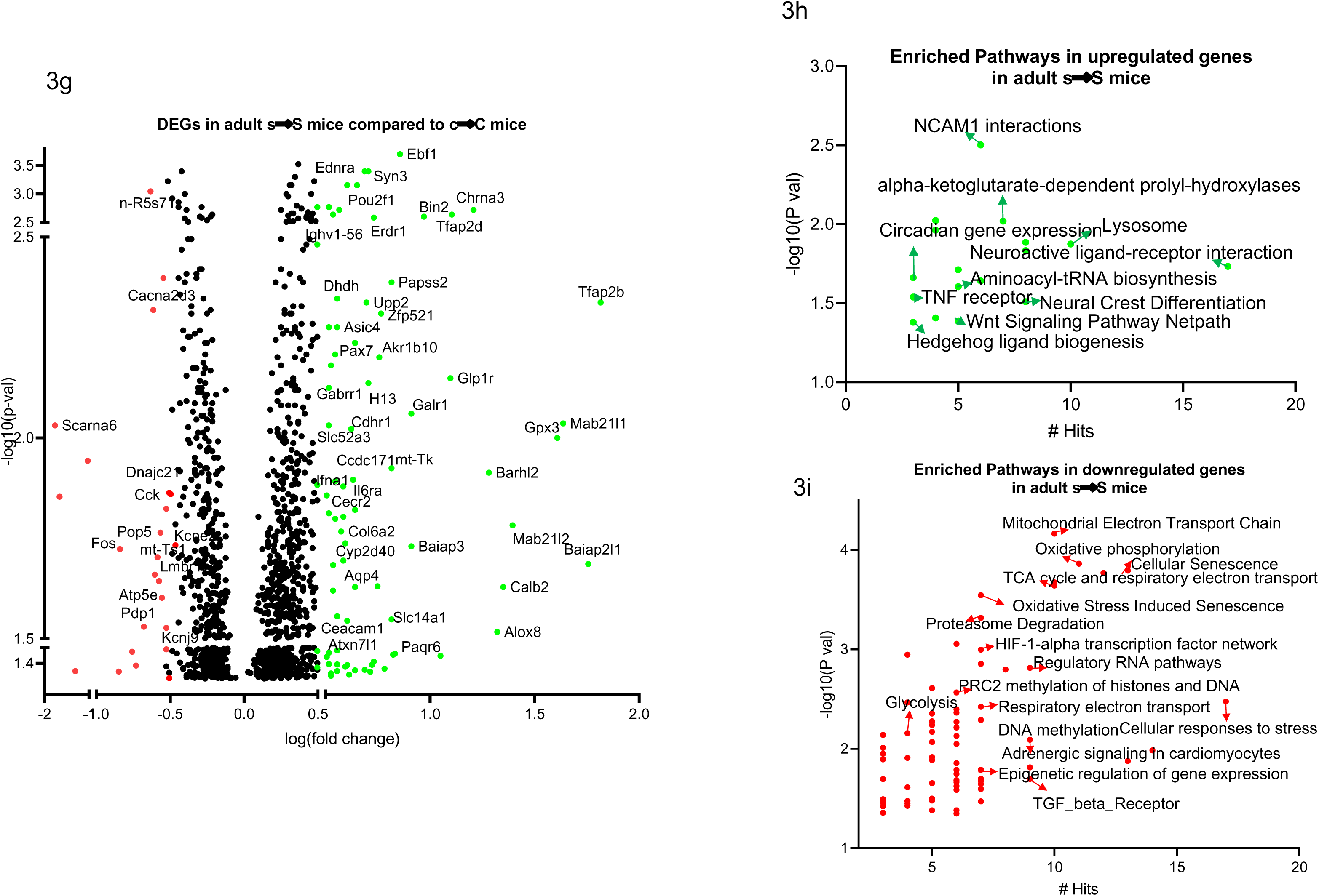
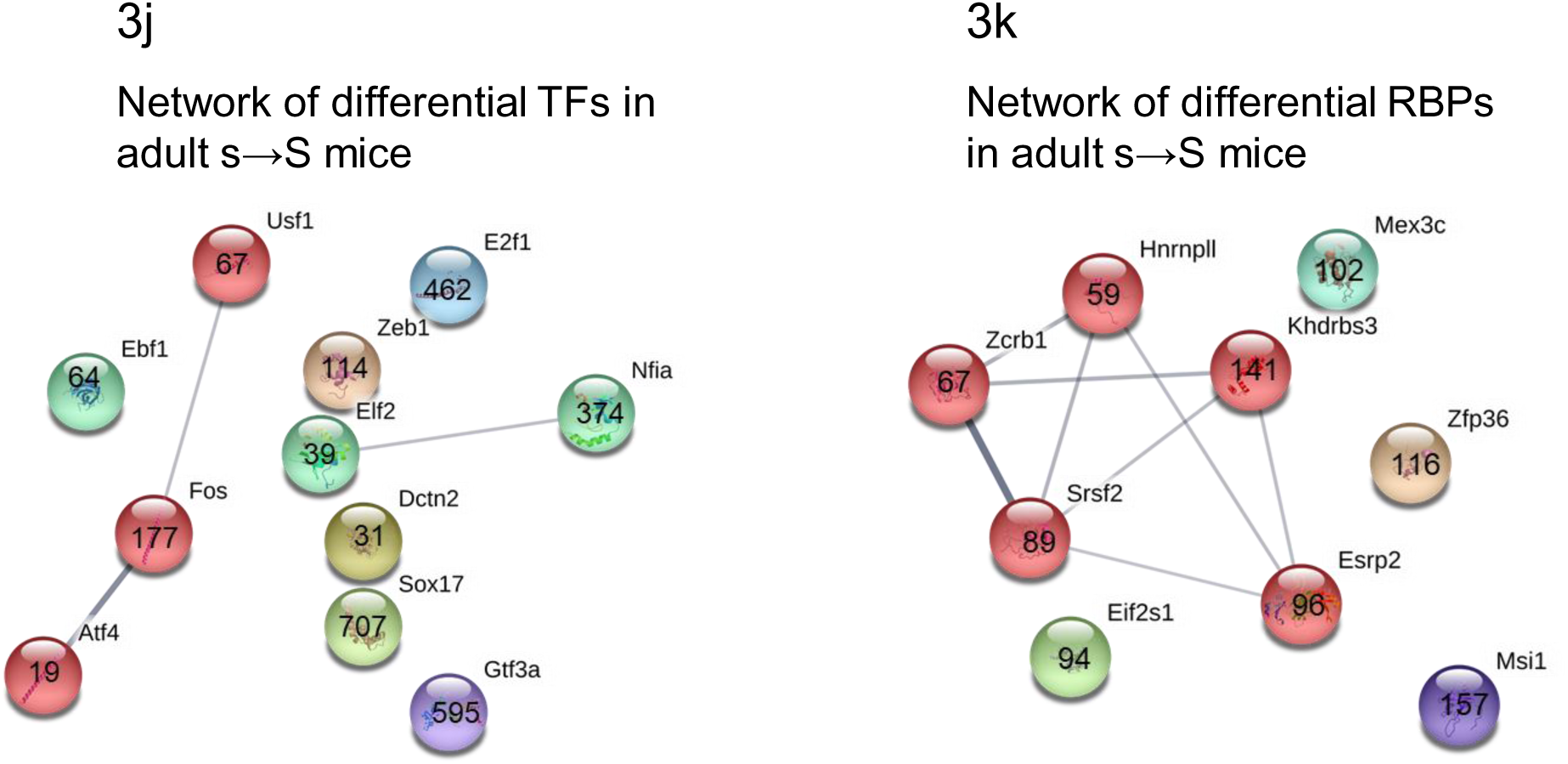
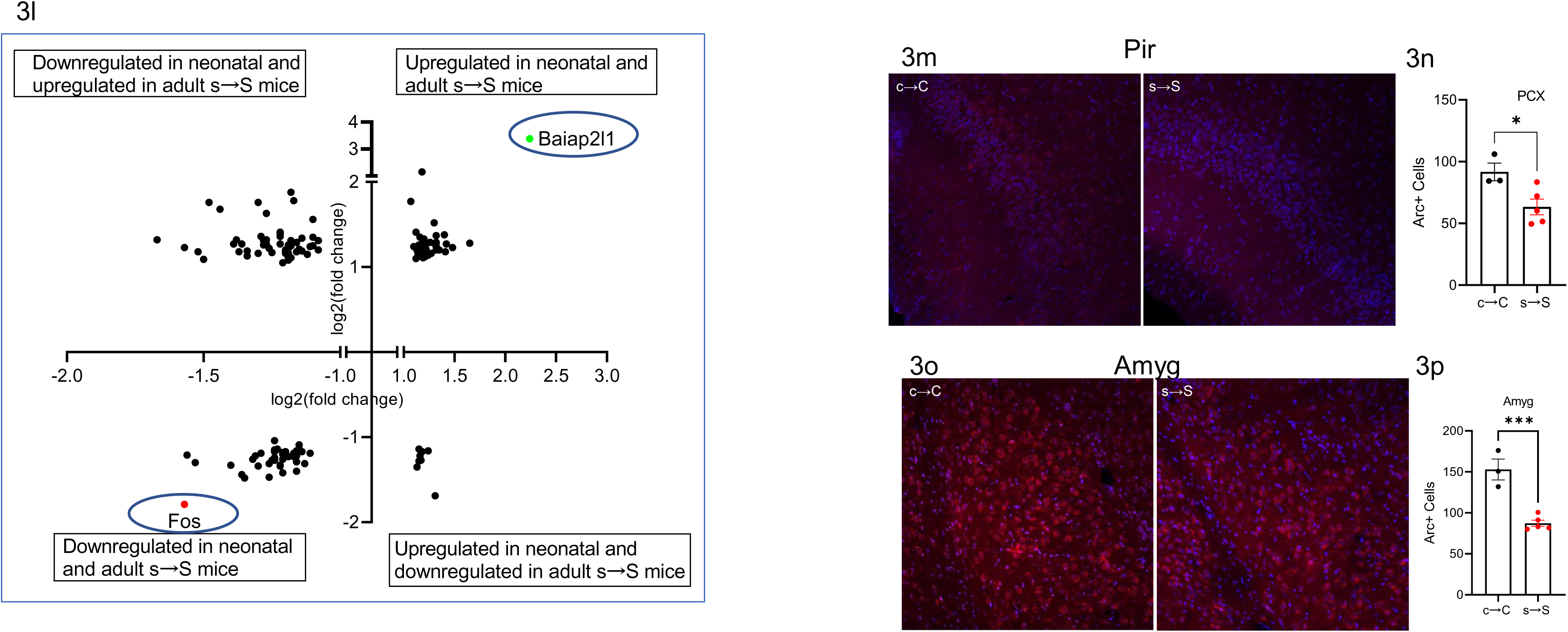
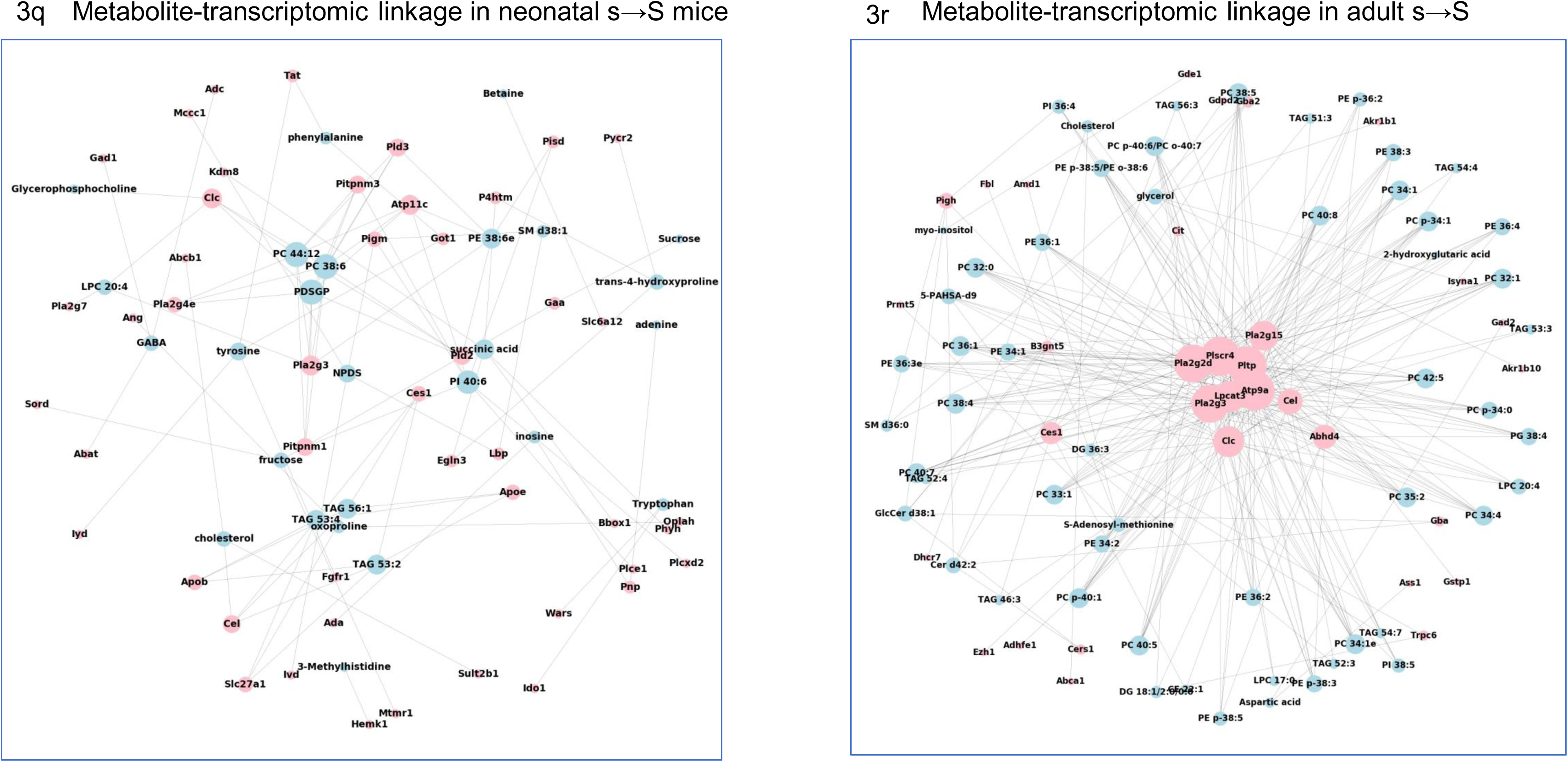
Prenatal exposure to stress causes alteration in brain transcriptomic signature. **a.** Volcano plot showing differentially expressed genes in brains from neonatal s→S versus c→C mice. Baiap2l1 exhibited the largest expression change among the upregulated genes (green symbols, log(fold) >0.5); Thy1 exhibited the largest expression change among the downregulated genes (red symbols log(fold) < -0.5). **b.** Real-time qPCR validation of microarray in neonatal mice. Relative global mRNA levels of Egr1, Fos, Avp, Vgf, Mcm6, and C1ql1 genes in the mice brain assessed by quantitative real-time PCR (n = 5). Unpaired student test (Egr1: t=6.346, *P*=0.0002; Fos: t=4.027, *P*=0.0038; Avp: t=6.807, *P*=0.0001; Vgf: t=5.172, *P*=0.0009, Mcm6: t=5.950, *P*=0.0003; C1q11: t=2.563, *P*=0.0335): neonatal c→C vs neonatal s→S, \**P*<0.05, \*\**P*<0.01, \*\*\**P*<0.01. Data are presented as means±S.E.M. **c,d** Significantly enriched biological pathways in the upregulated **(c)** and downregulated **(d)** genes in the neonatal mice with number of genes’ hits. Hypergeometric test, *P*<0.05. **e,f.** Differential **(e)** TFs and **(f)** RBPs, whose target gens are DEGs in the neonatal mice. STRING network visualization, using Markov clustering algorithm (MCL), with the number of their differential target genes. **g.** Volcano plot showing differentially expressed genes (DEGs) in brains from adult s→S versus c→C mice. **h,i.** Significantly enriched biological pathways in the upregulated **(h)** and downregulated **(i)** genes in the adult mice with number of genes’ hits. Hypergeometric test, *P*<0.05. **j,k.** Differential **(j)** TFs and **(k)** RBPs whose target gens are DEGs in adult s→S mice. STRING network visualization, using Markov clustering algorithm (MCL), with the number of their differential target genes. **l.** Volcano plot of genes showing similar and opposite directions of change in neonatal and adult s→S pups **m-p**. Immunohistochemistry analysis of Arc in the brains of adult s→S and c→C mice; **m,o.** Representative images of Arc-immunoreactivity in the **(m)** PCX, **(o)** Amyg, **n,p.** quantification of Arc immuno-positive cells in the **(n)** PCX, **(p)** Amyg. Unpaired t-test, PCX: t=2.83, *P=*0.029, n=3 C 5 S; Amyg: t=6.165, *P=*0.0008, n=3 C 5 S; Scale bar = 20uM. Values represent mean ± SEM. piriform cortex (PCX), and amygdala (Amyg), **q,r.** Network graphs were drawn to visualize the metabolite-transcriptome interaction in **(q)** neonatal s→S pups and **(r)** adult s→S pups, using “spring” layout from the NetworkX package with Python. Each blue node presents a metabolite, and each red node presents the enzyme-coding gene. If a gene was involved in the metabolic reaction of a metabolite, an edge was drawn to connect these two nodes. The size of the node was proportional to its connectedness in the graph: the more connected this node is, the bigger its size.

Pathway analysis using Hypergeometric test and Pathway Common Database and the ConsensusPathDB in neonatal s→S mice revealed the enrichment of mRNA processing, extracellular matrix organization, mitochondrial transport, GPCR signaling, glutamate transmission, glycolytic processes, ATP production, sympathetic system activation, and circadian regulation of gene expression (Fig. 3c,d, Table S4). After correction for multiple testing (FDR q < 0.05), mRNA processing remained enriched (Table S4).

Aligned with the gene regulation enriched pathways, 57 transcription factors (TFs) and 13 RNA binding proteins (RBPs) were differentially expressed in the s→S mice (Table S5). Bioinformatic analyses using MotifMap and ChiPseq databases reveal that many of the DEGs in the neonatal s→S mice are indeed targets (differential target genes (DTGs)) for more than one of the differential TFs and RBPs (Fig. 3e,f, Fig. S2a-h, Table S5).

In the adult s→S mice, the top downregulated genes were Fos, Cacna2d3, Zfp874b, mt-Ts1, several small nucleolar RNAs, and ribosomal proteins (Fig. 3g, Table S3). Among the top upregulated genes were mitochondrial genes and genes known to regulate energy metabolism: Glp1r, Tfap2b/d, Gpx3, Calb2, Alox8, Chrna3, Galr1, Ebf1, Paqr6, Gdpd2, Akr1b10, Syn3, Aqp4, Il6ra, and Tacr3 (Fig. 3g, Table S3).

The hypergeometric test identified enrichment of pathways that are associated with cellular responses to stress, oxidative phosphorylation, Electron Transport Chain, glycolysis, TCA cycle and respiratory electron transport, acyl-carnitine pathways, HIF-1-alpha transcription factor network, oxidative stress induced senescence, circadian gene expression, DNA and histone methylation, sympathetic signaling in cardiovascular system, and epigenetic regulation of gene expression (*P* < 0.05, Fig. 3h,i, Table S4). After correction for multiple testing (FDR q < 0.05), mitochondrial Electron Transport Chain remained enriched (Table S4).

Aligned with the gene regulation enriched pathways, 52 TFs (40 upregulated and 12 downregulated), and nine RBPs were differentially expressed in the brains of the adult s→S mice (Fig. 3j,k, Table S4). Several differential genes were identified as targets for these differential TFs and RBPs (Fig. 3j,k, S3a-h, Table S4).

Eighty-three genes followed the same change direction in neonatal and adult s→S mice, whereas 62 genes exhibited opposite alterations (Fig. 3l). The genes exhibiting the greatest degree of differential expression that changed in the same direction in neonatal and adult s→S mice were Baiap2l1 and Fos (Fig. 3l).

Given the alteration in many genes involved in synaptic plasticity and glutamate transmission in both neonatal and adult s→S mice, we analyzed relative expression of activity-regulated cytoskeleton-associated protein (Arc/Arg3.1), a critical gene for proper glutamatergic synaptic function and plasticity [59, 60]. We found that in the s→S adult pups, there were lower numbers of Arc-positive cells in the amygdala, hippocampus, striatum, and piriform cortex, with no change in the frontal cortex and nucleus accumbens, compared to c→C adult pups (Fig. 3m-p, Fig. S4a-h), though the total numbers of cells in these regions were not altered in s→S group, evidenced by the similar number of DAPI-stained nuclei.

### Bioinformatic integration of transcriptomic and metabolomic data substantiates mitochondrial and epigenetic mechanisms

Our metabolic profiling was in good agreement with gene expression data. The activities of the metabolites and their corresponding genes involved in the metabolic reactions were linked together through known enzymatic reactions from HMDB. The linkage analysis of the changes in gene expression levels with the changes in metabolite levels in the neonatal pups confirm the master biological pathways including mRNA modification, response to stress and hypoxia, oxidative-reduction process, phospholipid metabolic process, and DNA demethylation. The molecular functions enrichment analysis identified enzymatic activities of 2-ketoglutarate-dependent dioxygenase, phospholipase, and cytidine deaminase (Fig 3q,r, Table S6). The linkage analysis of the DEGs with the changes in metabolite levels in the adult pups revealed master pathways including chromatin modification, lipid metabolism, and glutamate pathways. Enriched chromatin modification (histone methylation (H4-R3 and H3-K27) was associated with enrichment in histone methyltransferase activity. Enriched lipid metabolism was linked with phospholipase activity, and phospholipid transporter activity (transferring acyl group) (Fig. 3q,r, Table S6).

### Distinct behavioral and Transcriptomic impacts of in utero exposure vs. impaired maternal care

Since mothers exposed to stress during pregnancy exhibited impaired maternal care and increased depressive-like behavior, we applied cross-fostering procedures to examine the distinctive effects of the *in utero* exposure vs. impaired maternal care upon the behavioral phenotype in adulthood (Fig. 4a). We found that pups born to control mothers and reared by stressed mothers (c→S mice) displayed similar behavioral phenotypes to those born to and raised by their stressed biological mothers: the c→S mice displayed social deficits (*P<*0.001, two-way ANOVA followed by Bonferroni post-test, Fig. 4b), and increased immobility time in the forced swim test (*P<*0.01, one-way ANOVA, followed by Tukey post-test, Fig. 4c). These results support a crucial role for the early life environment in the observed phenotypes. However, the caregiving by control females of mice born to stressed females (s→C) did not reverse the behavioral deficits. Thus, although s→C mice spent more time with the unfamiliar mice than the empty cup, the difference in time spent was not statistically significant (Fig. 4b). Similarly, caregiving by control mothers slightly reduced immobility time of s→C mice in the forced swim test; however, the levels were not significantly different from those of s→S mice (*P>*0.05, when compared to c→C or s→S mice, Fig. 4c).

**Figure 4.**
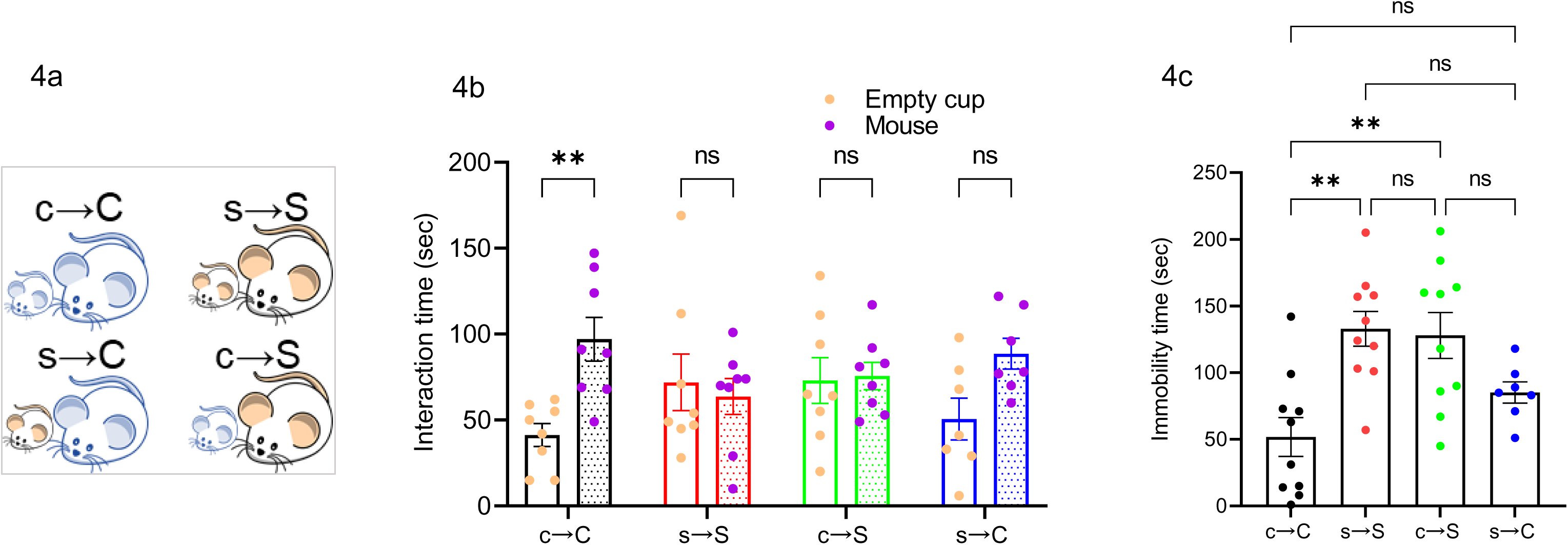
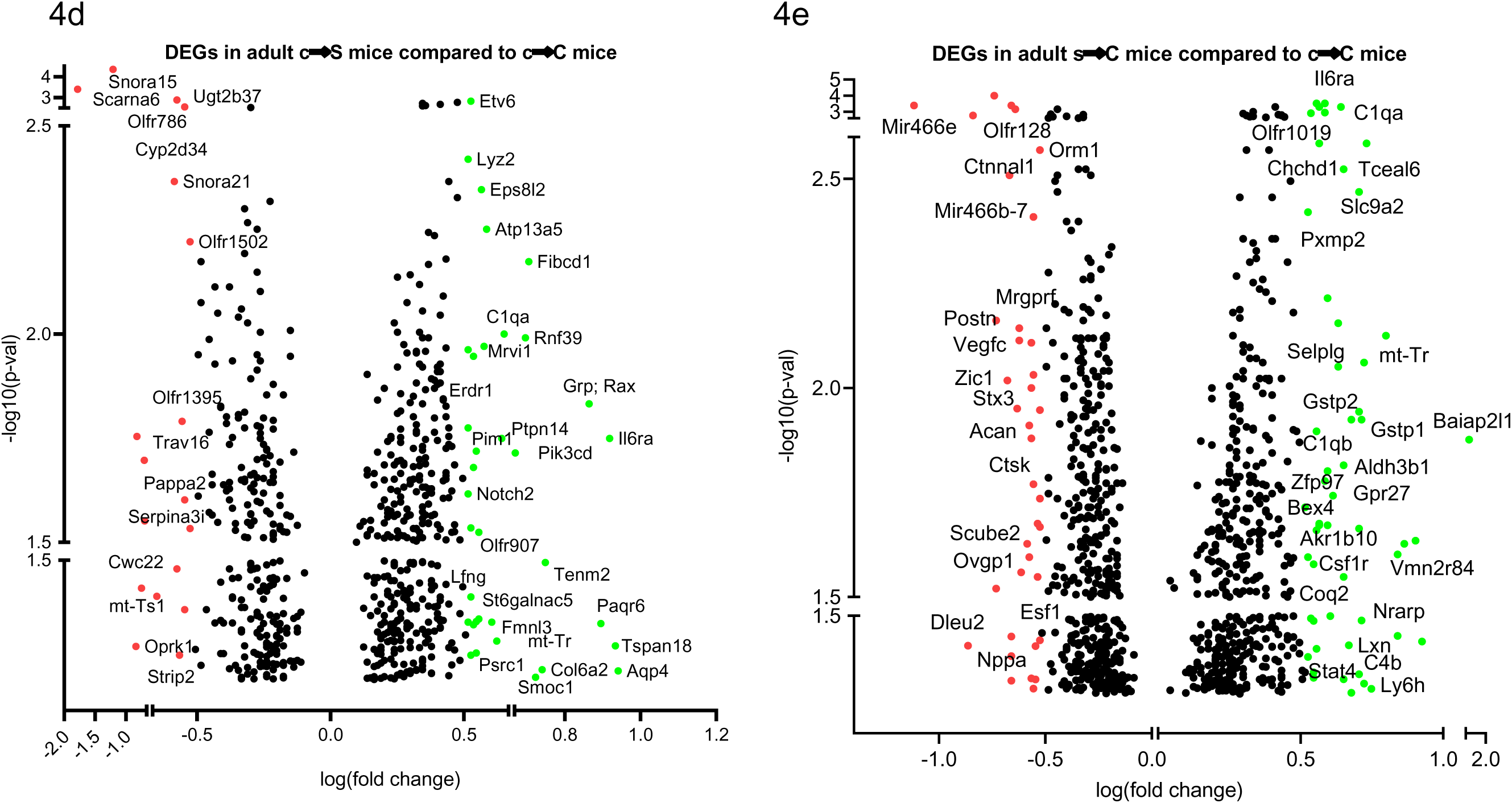
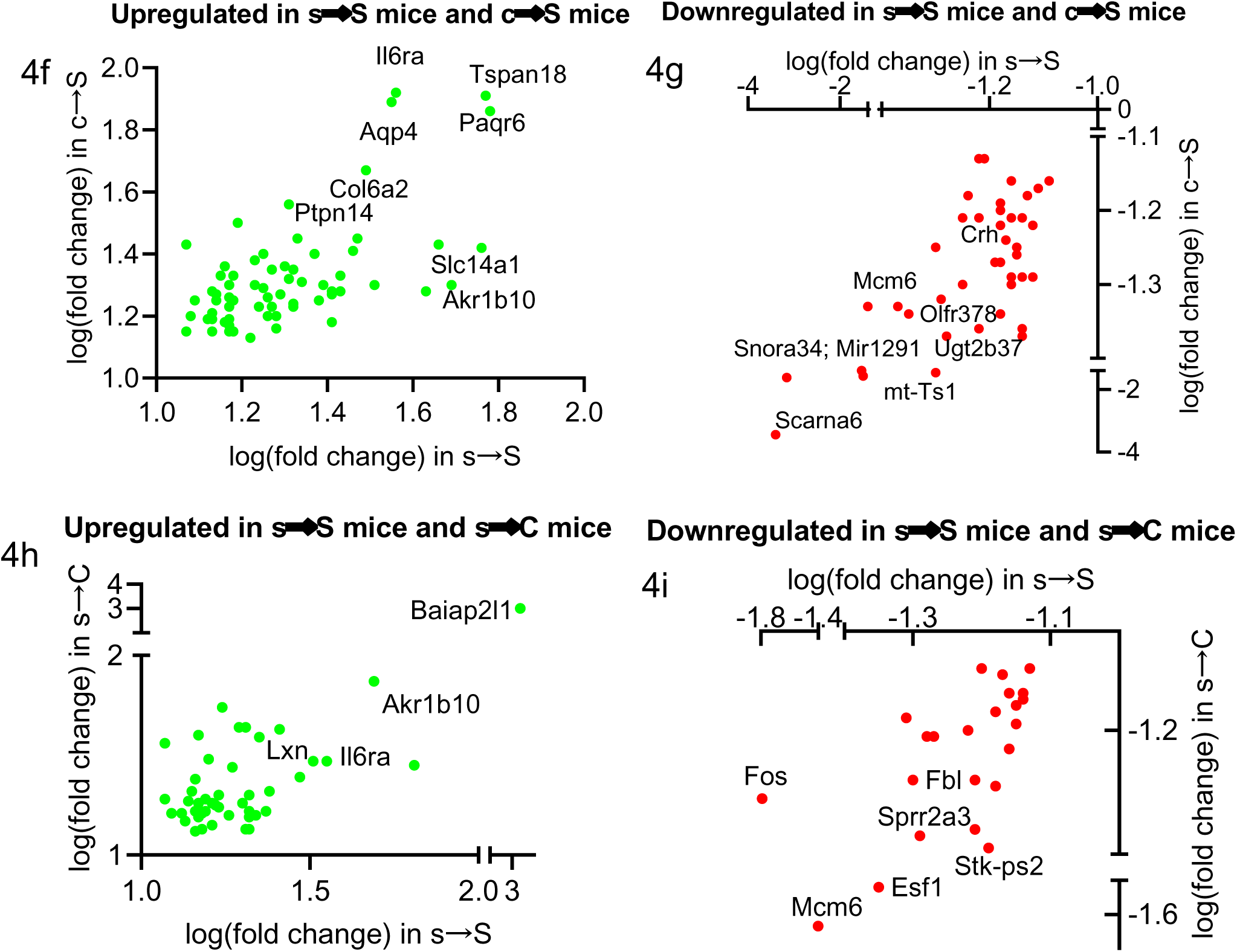
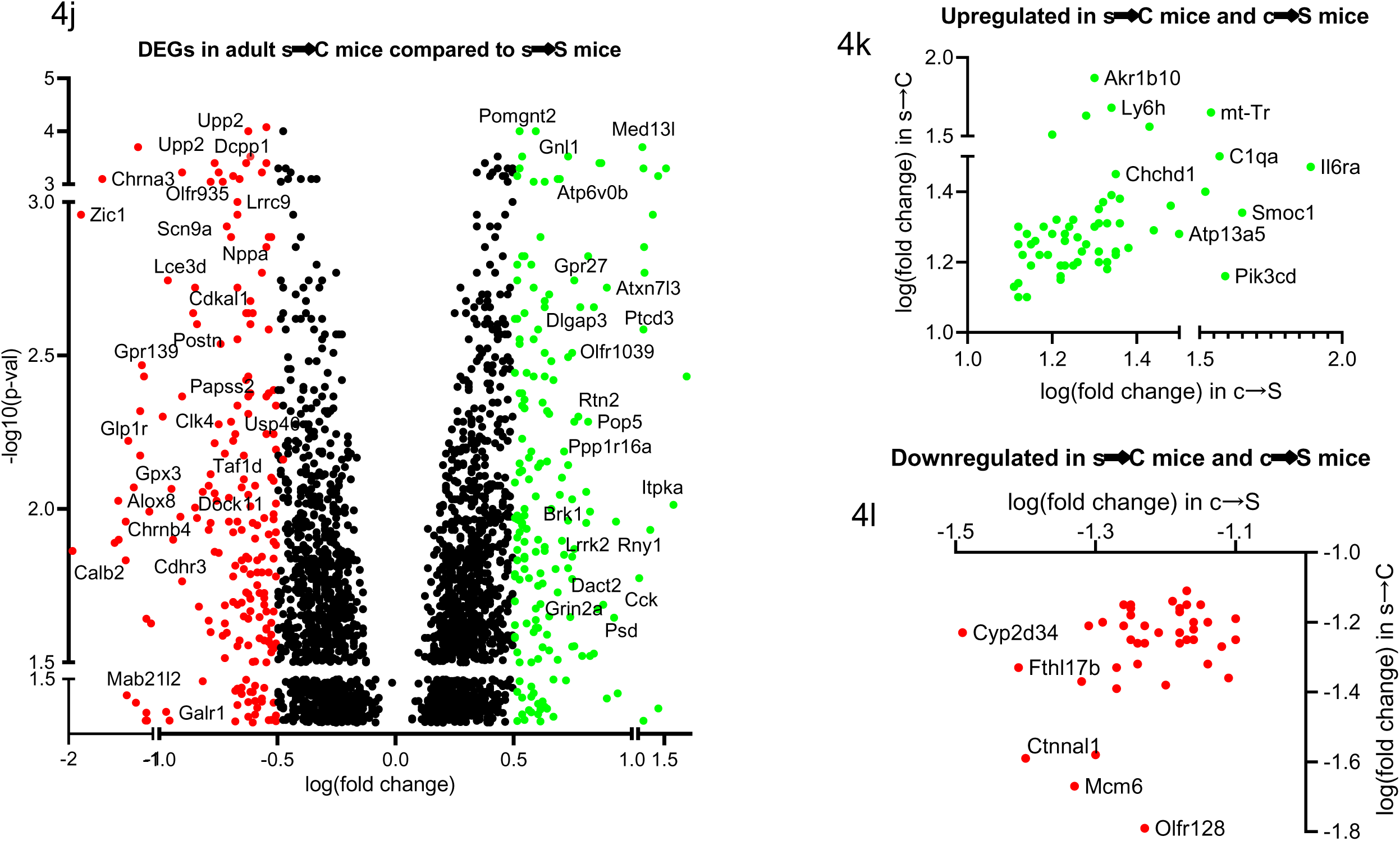
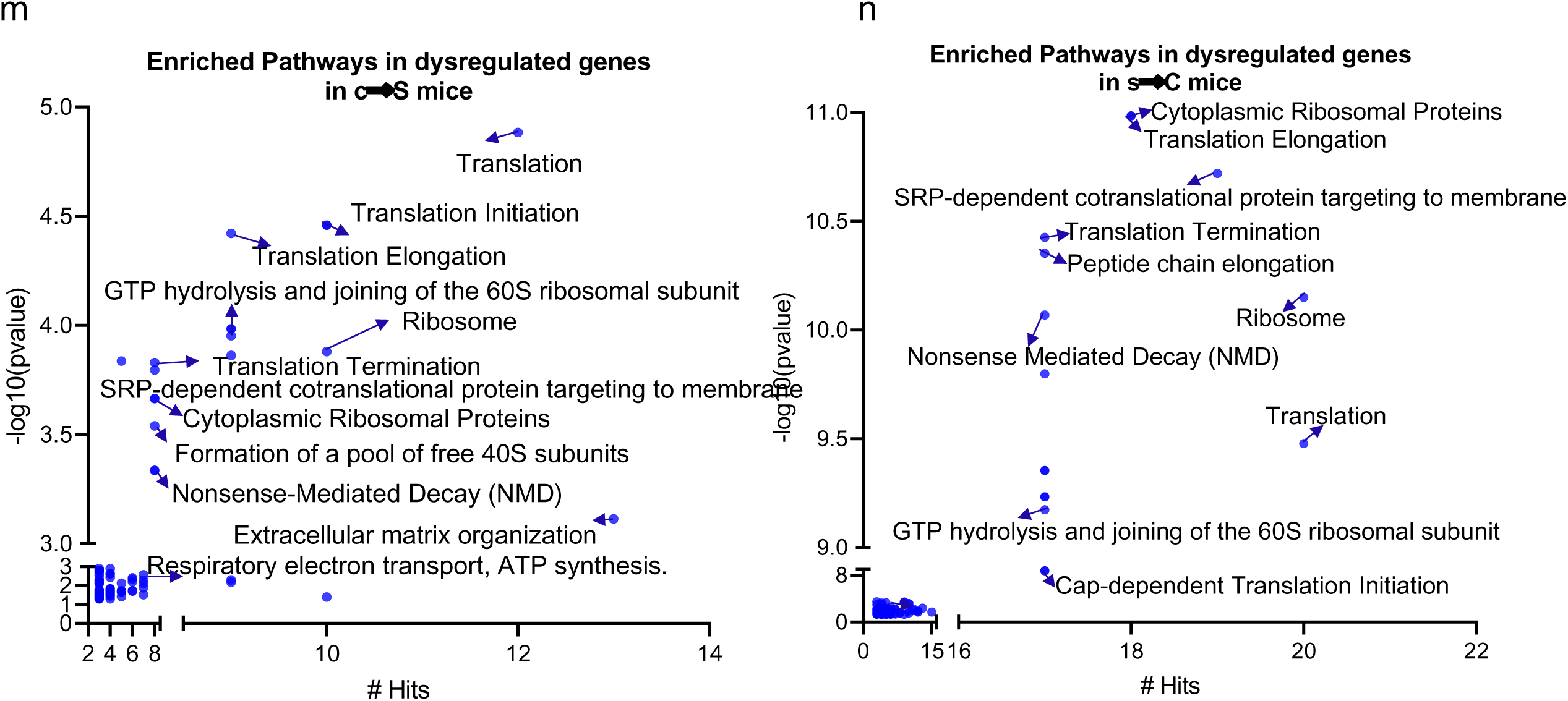
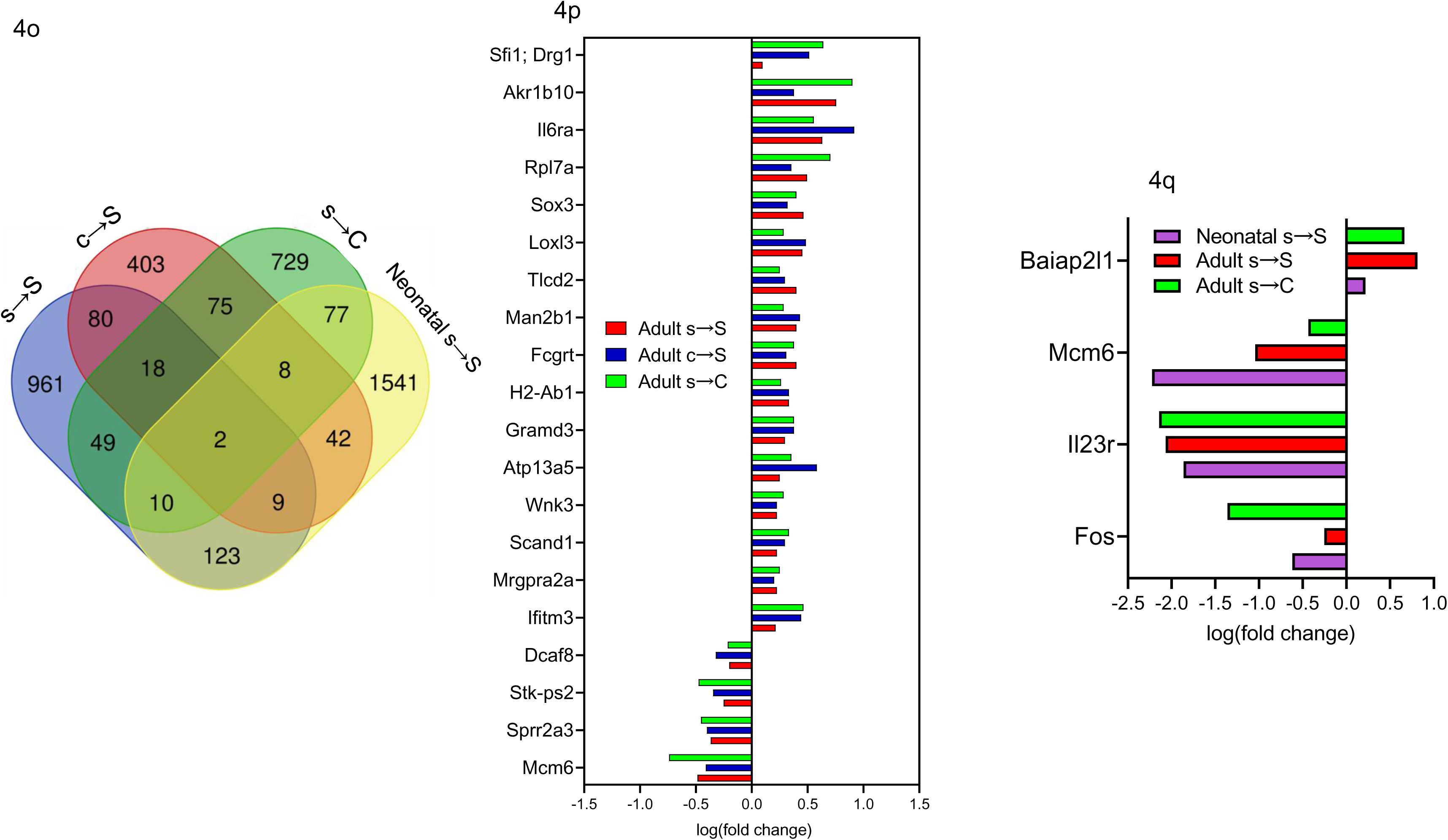
Cross-fostering effects on behavioral phenotype and transcriptomic profile. **a.** Schematic of the Cross-fostering experiment design **b.** Time mice spent interacting with empty cup and unfamiliar mouse in the social interaction assay (n=8 c→C, 8 s→S, 8 c→S, 7 s→C). Two-way ANOVA: object effect (F_1,54_=7.255, *P*=0.0094), exposure to stress effect (F_3,54_=0.1260, P=0.9443), stress x object interaction (F_3,54_=3.422, *P*=0.0235) followed by Bonferroni post-hoc test: empty cup vs unfamiliar mouse, \*\**P<*0.01, ns: not significant. **c.** Time mice spent immobile in the forced swim assay (n=10 c→C, 10 s→S, 10 c→S, 7 s→C). One way ANOVA revealed a significant stress effect (F_3, 33_=7.531, *P*=0.0006) followed by Tukey post-hoc test: c→C vs s→S vs c→S, \*\**P<*0.01, ns: not significant. **d.** Volcano plot showing DEGs in brains from adult c→S versus c→C mice. **e.** Volcano plot showing DEGs in brains from adult s→C versus c→C mice. **f,g.** Volcano plot of genes showing similar direction of change in c→S and s→S: **(f)** upregulated and **(g)** downregulated genes. **h,i.** Volcano plot of genes showing similar direction of change in s→C and s→S: **(h)** upregulated and **(i)** downregulated genes. **j.** Volcano plot showing DEGs in brains from adult s→C versus s→S mice. **K,l.** Volcano plot of genes showing similar direction of change in c→S and s→C mice: **(k)** downregulated and **(l)** upregulated genes. **m,n.** Enriched biological pathways in the DEGs of **(g)** c→S and **(h)** s→C mice with number of genes’ hits. Hypergeometric test, *P* < 0.05. **o.** Venn diagram showing the overlap of DEGs in all mouse groups. **p.** Overlap of DEGs in adult mouse groups that received one hit of stress, prenatally only (s→C), postnatally only (c→S) or two hits of stress (s→S). **q.** Overlap of DEGs in mouse groups that received at least one hit of stress prenatally.

### Distinct Transcriptomic impacts of in utero exposure vs. impaired maternal care

Transcriptomic analysis in adult c→S and s→C mice revealed 640 and 973 DEGs, respectively, compared to c→C mice, of which 108 and 69 genes in adult c→S and s→C mice, respectively, were altered in the same direction of the s→S DEGs (Fig. 4d,e, Table S3).

Among the top DEGs that displayed the same direction of change in both s→S and c→S mice were Il6ra, Aqp4, Atp13a5, Akr1b10, Tspan18, and Paqr6 (upregulated) (Fig. 4f), and Scarna6, Snora34, Mcm6, Akr1c20, Mir1291, and Crh, (downregulated) (Fig. 4g). On the other hand, among the top DEGs that displayed similar changes in s→S and s→C mice were Il6ra, Baiap2l1, Akr1b10, Gstp1, and Lxn (upregulated) and Fos, Mcm6, Fbl, Stk-ps2, and Esf1 (Fig. 4h,i). Most interestingly, caregiving by control mothers reversed the alterations in 559 genes (Fig. 4j). Among the top DEGs in s→C group compared to s→S group are Gpx3, Galr1, Chrna3, Alox8, Calb2, Glp1r, Tfap2d, and Mab21l2, which were downregulated, and Atp5e, Crhbp, Cck, and Pop5, which were upregulated (Fig. 4j).

Mice exposed to one hit of stress either prenatally (s→C mice) or postnatally (c→S mice) shared 103 DEGs, which exhibited changes in the same directions in both s→C and c→S mice (Fig. 4k,l). Hypergeometric distribution and Fisher’s exact test revealed common significant enriched pathways between s→C and c→S mice, including mRNA translation (initiation, elongation, and termination) and mRNA decay (FDR q < 0.05, Fig. 4m,n, Table S4).

Further, 20 DEGs overlapped across the three adult groups that were exposed to one hit (s→C and c→S mice) or two hits (s→S) of stress (Fig. 4o,p). When considering the neonatal transcriptomic changes, we found only four genes that exhibited early- and long-lasting expression changes in the animals that were exposed at least prenatally to stress (neonatal s→S, s→S, and s→C) (Fig. 4q). Only one gene (Mcm6) displayed the same change in all animals that were exposed to stress prenatally and/or postnatally and was changed as early as 24-hr after birth.

### Transcriptomic-overlapping between human MDD and the four stress mouse groups

The transcriptomics linkage analysis of the DEGs in the four groups of S mice with gene expression changes in human major depressive disorder (MDD) revealed 203, 137, 105, and 74 DEGs in the neonatal s→S, adult s→S, s→C, and c→S mice respectively that overlap with human MDD (Fig. 5, Table S7). Fisher exact test revealed significant overlaps between human MDD and all the stress mouse groups, with *P* values of 1.33E-31, 3.34E-21, 1.09E-19, and 4.58E-11 for intersections between MDD and neonatal s→S, adult s→S, s→C, and c→S respectively.

**Figure 5.**
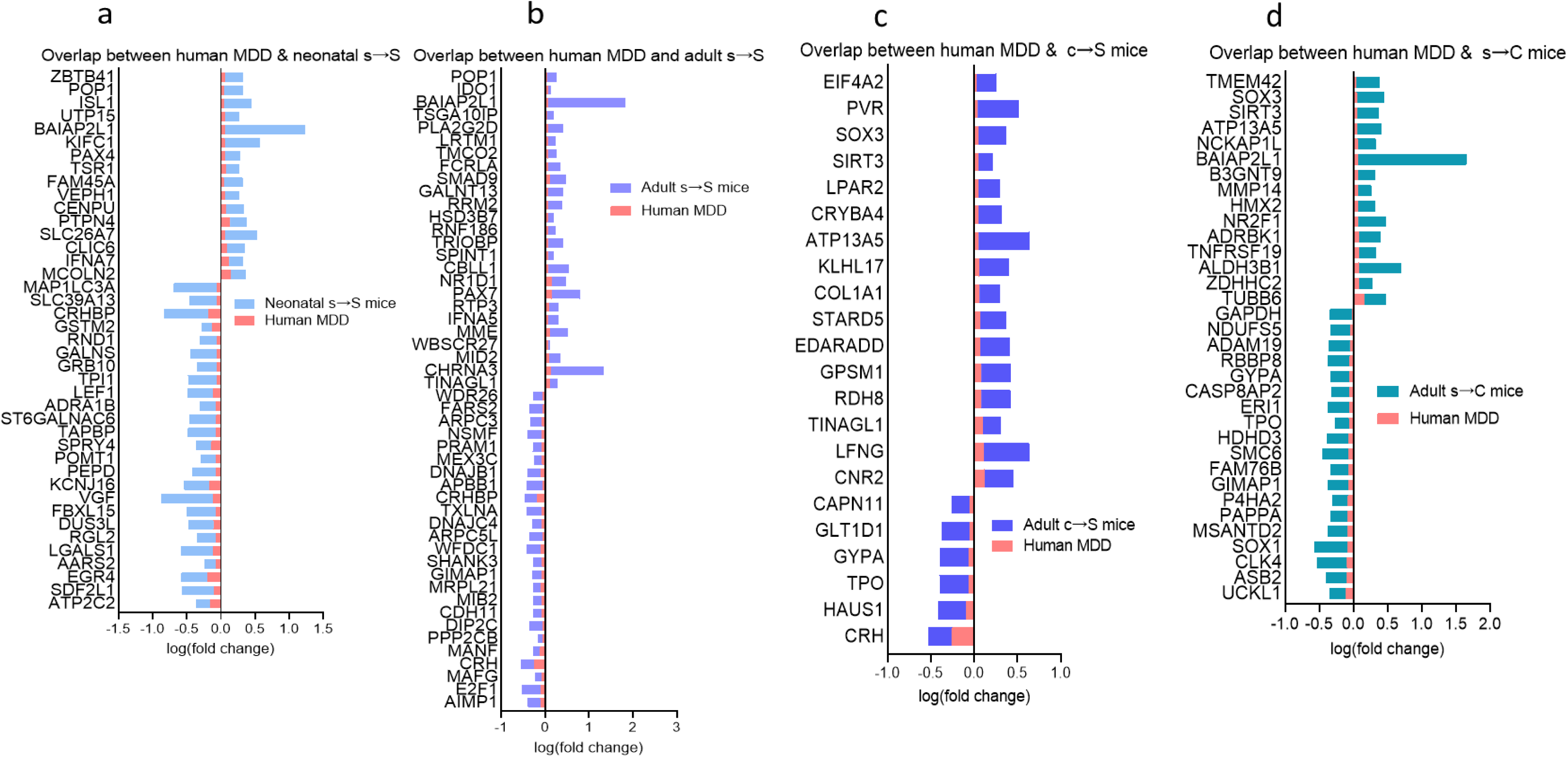
Integrated mouse-human transcriptomic analysis reveals overlap between genes whose expressions changed in mice exposed to stress and MDD patients. **a.** Top common DEGs and their log(fold change) in MDD patients and 24-hr s→S mice. **b.** Top common DEGs and their log(fold change) in MDD patients and adult s→S mice. **c.** Top common DEGs and their log(fold change) in MDD patients and c→S mice. **d.** Top common DEGs and their log(fold change) in MDD patients and s→C mice.

### Acetyl-L-carnitine (ALCAR) protected against and reversed depressive-like behavior induced by prenatal trauma

Given that the behavioral deficits in the s→S mice were associated with a decrease in palmitoyl-carnitine, increase in 2-HG, and impairments in the mitochondrial functions, lipid β-oxidation and phospholipid metabolism, we tested whether the behavioral deficits could be rescued by an agent, acetyl-L-carnitine (ALCAR), that is known to restore these functions. ALCAR is known to elicit its antidepressant effects through a number of mechanisms including rescuing mitochondrial functions and lipid β-oxidation metabolism, improving glutamate transmission, and enhancing chromatin acetylation by providing an acetyl group to chromatin [61, 62]. ALCAR was supplemented in the drinking water of s→S mice either from weaning to adulthood (3-8 weeks), or for one week in adulthood (7-8 weeks) (Fig. 6a). ALCAR supplementation for one week during adulthood rescued the depressive-like behavior (*P<*0.001, Fig. 6b) in s→S mice. One week after ALCAR cessation, however, the anti-depressant effect of ALCAR was diminished (Fig. 6c). ALCAR supplementation from weaning rendered s→S mice resistant to developing depressive-like behavior (*P<*0.01, Fig. 6b). Strikingly, ALCAR antidepressant effects lasted one week after its cessation (*P<*0.001, Fig. 6c).

**Figure 6.**
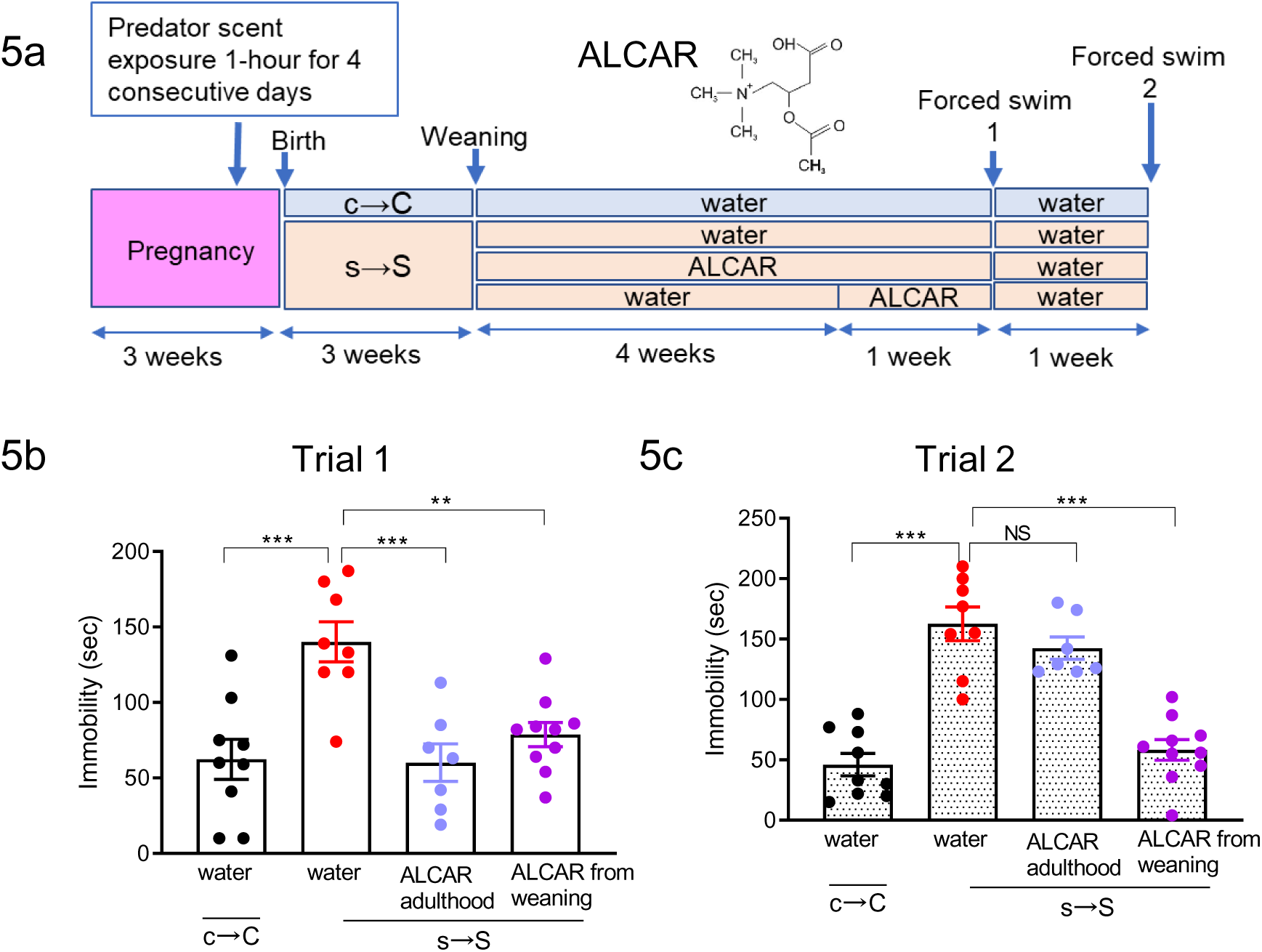
Acetyl-L-carnitine (ALCAR) reverses depressive-like behavior after its cessation. **a.** Schematic of the two ALCAR administration regimes, and the chemical structure of ALCAR **b.** Time mice spent immobile in the forced swim assay following the treatment of ALCAR for five weeks from weaning, and one week during the adult stage, (n= 9 c→C, 8 s→S, 7 s→S + ALCAR from Weaning, 10 s→S + ALCAR in Adulthood). One-way ANOVA: treatment effect (F_3,30_=9.72, *P=*0.0001) followed by Tukey post-hoc test, \*\**P<*0.01, \*\*\**P<*0.001, ns, not significant. **c.** Time mice spent immobile in the forced swim assay one week after the cessation of ALCAR treatment, One-way ANOVA revealed a significant treatment effect (F_3,30_=32, *P=*0.0001) followed by Tukey post-hoc test, \*\*\**P<*0.001, ns, not significant.

## Discussion

The long-term devastating impacts of prenatal traumatic stress are well established in humans and animals [1, 3, 4, 63]. Yet, the mechanistic involvement of prenatal stress independent of stress effects on maternal care is largely unknown.

Here, we present results demonstrating that exposure to trauma during pregnancy induces in the offspring long-lasting depressive-like behavior and social deficits, without affecting cognitive functions. Using cross-fostering experiments, we demonstrate that these behavioral deficits are associated with divergent and convergent mechanisms of both *in-utero* and early-life parenting environments. Thus, the exposure to postnatal early-life hit of stress, through raising of normal pups by traumatized mothers, produces a behavioral phenotype that is similar to that induced by a double-hit of stress (in pups born and raised by their biological traumatized mothers). Good caregiving by normal mothers of pups that were exposed to prenatal hit of stress, however, did not completely prevent the manifestation of trauma-induced behavioral deficits. Thus, our data demonstrate distinctive behavioral impacts of one-hit exposure to stress (either prenatally through or postnatally) or two-hits stress (prenatal and postnatal exposures).

Associated with the distinct behavioral deficits we found, through metabolomic, transcriptomic and bioinformatic analyses, mechanisms that involve stress- and hypoxia-response energy metabolic pathways, especially mitochondrial ATP production. These acute responses seem to have resulted in long-lasting adaptations in glycolysis, homeostasis of energy lipids, and epigenetic processes pertaining to DNA and chromatin modifications, as evidenced by the disruptions of these pathways in the brains of adult mice. We therefore propose a model through which stress exposure during pregnancy induces, in progeny, early and long-lasting mitochondrial metabolism dysfunctions and epigenetic changes, associated with social behavioral deficits (Fig. 7a). According to our model, the striking increase in the mitochondrial metabolite and epigenetic modifier 2-hydroxyglutarate (2-HG) in the brains of neonatal mice, whose mothers were exposed to extreme stress, likely forms the first step of the consequent stress-response events.

**Figure 7.**
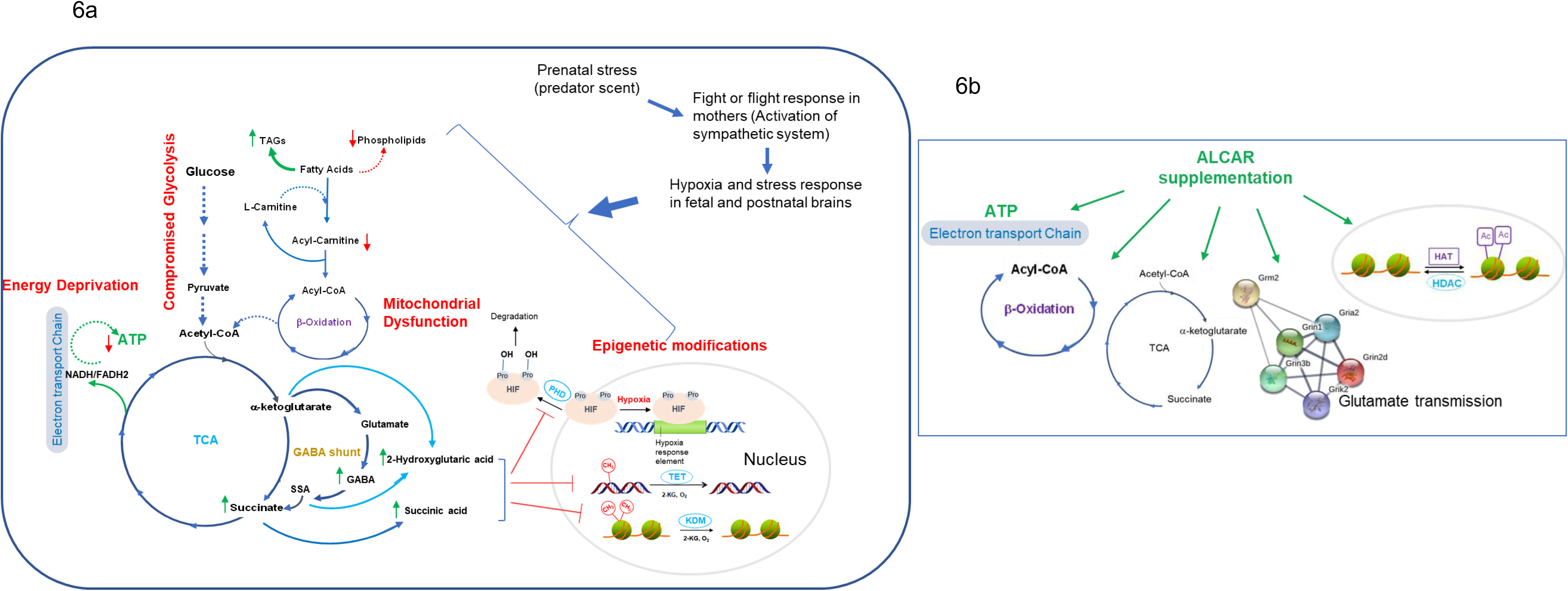
Model. **a.** Prenatal exposure to stress elicits depression through a number of mechanisms that may begin with hypoxia, which results in compromised glycolysis, energy deprivation, mitochondrial dysfunction (oxidative phosphorylation and ATP production), and altered epigenetic processes. **b.** Possible mechanisms through which ALCAR may produce its antidepressant effects.

Exposure to extreme stress during pregnancy is known to activate the maternal fight-or-flight sympathetic system, and produce fetal reduction of oxygen (hypoxia), which might be detected in the infants several months postnatally [64–68]. The mitochondrial metabolite 2-HG is an established marker for the hypoxia, defective mitochondria, decreased electron transport chain activity, and acidic cellular conditions [40, 41, 69, 70]. Accumulation of 2-HG in hypoxia occurs as a result of the TCA cycle dysfunction and, in turn, 2-HG inhibits electron transport and glycolysis to counterbalance the deleterious consequences of hypoxia [69]. Therefore, 2-HG accumulation in the neonatal s→S pups, accompanied by succinate accumulation and activated GABA shunt, indicates a hypoxic state and a disruption of the mitochondrial metabolism. The hypoxic state in newborn s→S mice seems to trigger adaptation mechanisms, particularly in metabolic and energy demand [71–73]. Accordingly, acute stress-hypoxia responses in the neonatal s→S mice affect glycolysis and shift mitochondrial-energy metabolism (disruption in ATP production, accumulation of TAGs and depletion of membrane phospholipids). In addition to its role in hypoxia, 2-HG is an epigenetic modifier, which regulate epigenetic programming through its competitive inhibition of 2-ketoglutarate (2-KG)-dependent enzymes [74]. 2-KG-dependent enzymes known to be inhibited by 2-HG include TET(1-3) DNA demethylase enzymes, Jumonji family of histone lysine demethylases (JmjC-KDMs), and prolyl hydroxylases (PHDs 1–3), which hydroxylate proline in hypoxia-inducible factors (HIFs), as a post-translational modification [75–79]. Through these mechanisms, 2-HG alters DNA and histone methylation [75, 76, 80, 81], and increase HIFs accumulation, resulting in selective upregulation of HIF-target genes [82, 83]. In support of HIFs accumulation, we found lower levels in the neonatal s→S pups of 4-hydroxyproline, a marker for the degradation of the proline-containing HIFs [84]. The reduced levels of 4-hydroxyproline in the neonatal s→S pups indicate decreased hydroxylation and, hence, lower degradation of HIFs, causing HIFs accumulation.

HIFs target genes include practically all genes of the glycolytic pathway [85–87]. While these metabolic and epigenetic changes aim to protect the brain from hypoxia during development, their effects may persist and confer the susceptibility for psychiatric disorders including depression later in life. First, the perturbation of the glycolysis pathways during stress triggers the activation of GABA shunt as an alternative pathway to enhance energy production [49, 88–90]. Ultimately, the stress-associated depletion of the brain’s energy resources leads to alteration in GABA and glutamate transmission and results in impaired neuronal plasticity underlying depression [91, 92], reflected by the decreased Arc expression in brain regions involved in emotion and stress. Second, the disruption of glycerolipids homeostasis exacerbate neural function and has acute and long-lasting impacts through directing the lipid flux toward energy storage rather than membrane extension or signaling. Lastly, while the early life 2-HG-triggered epigenetic modifications serve to offset mitochondrial dysfunction in response to stress [92], they may produce long lasting adaptations by altering methylation states and the expression of key genes involved in neurodevelopment, neuron maturation and differentiation, axon-genesis, and synaptic plasticity [93, 94].

Our findings indicate a two-hits mechanistic of convergence as well as divergence of some key metabolic, epigenetic and neuronal pathways induced by prenatal and early life trauma, which are involved in the pathophysiology of depression. This is supported by the overlap in specific sets of genes and pathway networks among s→S, s→C and c→S mice. Our finding of common genes in human depression and the four experimental animal groups (neonatal s→S, adult s→S, s→C and c→S) provides a translational aspect of our model and support for our convergent pathways hypothesis of the emergence of stress-induced depression. Strikingly, among the different stress mouse groups, the neonatal s→S mice exhibited the highest overlap in DEGs with human MDD. This finding strongly indicates the importance of our model in identifying novel biomarkers that serve as predictive markers in early life for the risk of the development of depression in adulthood, and in providing a potential target for early preventive approaches and therapeutic strategies.

Thus, a pharmacological intervention that restores the perturbed pathways at the appropriate time of life would be predicted to protect against depression. ALCAR possesses unique features that make it an ideal candidate, providing a rare opportunity for preventive rather than therapeutic intervention for depression. First, ALCAR has long been known to enhance mitochondrial function and facilitate ATP production [61, 62]. Second, ALCAR promotes transportation of fatty acids into the mitochondria for subsequent oxidation and ATP generation and, thus, corrects the lipid profile, and directs the flow of lipids toward membrane production and energy production and TAGs reduction [24, 95]. Third, ALCAR produces epigenetic modifications that involve histone acetylation (through providing acetyl group from ALCAR to chromatin) [61, 62]. Through its epigenetic actions, ALCAR regulates the expression of key genes crucial for synaptic plasticity [61, 62]. Fourth, ALCAR has been shown to correct glutamate transmission in animal models of stress [61, 62]. Lastly, and most importantly, several randomized clinical studies demonstrated the effectiveness of ALCAR supplementation to decrease depressive symptoms [96, 97]. Indeed, the antidepressant effects of ALCAR have been speculated to occur through more than one of these mechanisms including enhancing mitochondrial function and facilitate ATP production [61, 62].

We found that ALCAR antidepressant effects outlasted its treatment end when it was administered early in life (at weaning time), but not when it was administered in adulthood. The timing and duration of ALCAR administration might be deterministic factors in the long-lasting anti-depressant effect of ALCAR, given that animals administered ALCAR from weaning time received the drug for five weeks, whereas those administered ALCAR in postnatal week 7 received it for one week. However, considering that one week cessation of ALCAR is sufficient as a washout period, the long-lasting effect of ALCAR seems to be more likely due to the administration timing (weaning time vs adulthood) rather than duration (one week vs 5 weeks).

This long-lasting anti-depressant effect of ALCART after its cessation suggests epigenetic mechanisms, through which ALCART likely increases histone acetylation causing chromatin structure remodeling. Since prenatal stress induced in s→S mice changes in all pathways that can be improved by ALCAR (mitochondrial functions and lipid β-oxidation metabolism, glutamate transmission, and epigenetic modifications), and given the distinctive effects of one-week vs 5 weeks of ALCAR administration, it is likely that ALCAR elicited its antidepressant effects on s→S mice via more than one mechanism (Fig. 7b). These novel findings provide a unique opportunity for utilizing ALCAR as a prophylactic supplementation to young children and adolescents to protect against depression in high-risk populations. Remarkably, ALCAR levels are low in depression patients, and the declines are greater in patients with a history of childhood trauma and emotional neglect [98].

Given the unique features of ALCAR, this natural supplement can represent an innovative and unique prophylactic and therapeutic strategy, should it be administered at the appropriate time of life. This has the potential to change the lives of millions of people who suffer from major depression or have the risk of developing this disabling disorder, particularly those in which the depression arose from prenatal traumatic stress.

In conclusion, we demonstrate that intergenerational trauma induces social deficits and depressive-like behavior through divergent and convergent mechanisms of both in utero and early-life parenting environments. We establish 2-HG as an early predictive biomarker for trauma-induced behavioral deficits and demonstrate that early pharmacological correction of mitochondria metabolism dysfunction by ALCAR can permanently reverse the behavioral deficits.

## Acknowledgements

The authors would like to thank Dr. Olivier Civelli for helpful discussions. The work of AA was supported by the department of pharmaceutical sciences-UCI. The work of SC and PB was supported in part by NIH grant GM123558. The work of GWA was supported by NIGMS grant GM130377.

## Author contributions

AA designed the experiments and wrote the manuscript; LM helped with designing the experiments; SA, AP, ZW, CP, CH, MK, and Zich W conducted behavioral experiments on the pups; LA conducted behavioral experiments on the mothers; EC and HB conducted the immunostaining experiments, AS and GA conducted the transcriptomics analysis, SC and PB conducted the bioinformatic analysis.

