## Supplemental Figures for "Intergenerational Stress Transmission is Associated with Brain Metabotranscriptome Remodeling and Mitochondrial Dysfunction"

***Corresponding Authors**

Dr. Amal Alachkar

Department of Pharmaceutical Sciences

University of California, Irvine, CA 92697

### Supplementary Methods

#### *Maternal behavior: pups' retrieval assay with postpartum mice*

Maternal behavior was examined in predator-odor-exposed dams and control dams using pups' retrieval assay. The assay was conducted on PPD2, in which, a maximum of five minutes of video recording time to calculate the mother's latency and duration. The mother was temporarily removed from the cage, and pups were removed except for three pups, which were placed on each of the three corners of the cage (not the nest corner). The mother was then returned to her nest and the latency and duration of pups' retrieval were recorded.

### Supplementary Results

*Mothers exposed to predator scent exhibit deficits in maternal behavior, and depressive-like behavior*

Pregnant mice were exposed to predator scent (PS) for one hour daily for four consecutive days from gestational day 17. The exposure to the PS triggered fear-like responses in the mice, displayed by escape behavior (avoidance) and heart palpitations.

Pup retrieval is a key indicator of maternal care. At postpartum day 2 (PPD2), stressed mothers (S) retrieved pups with significantly higher latencies (time to retrieve the first pup) and longer retrieval duration (retrieval of three pups) compared to the control mothers (C) ($P<0.05$ and $P<0.001$, for latency and duration respectively, Fig. S1a,b). In the forced swim test on PPD5, the S mothers exhibited higher immobility time than C mothers ($P<0.001$, Fig. 1c), indicating increased depressive-like behavior in these mice. Other behaviors of mothers were normal (Fig. S1d-g).

### Supplementary Figures

**Figure S1. Prenatal exposure to stress produces impairments in maternal behavior and depressive-like behavior in mothers**

**a-d.** Exposure to stress produces maternal behavior deficits

**a,b.** Pups retrieval latency and duration: **(a)** Time to retrieve one pup (n=13 c→C, 12 s→S), Mann Whitney test (U=24.50, *P*=0.0025): c→C vs s→S, ** *P*<0.01. (**b)** Time to retrieve all pups (n=13 c→C, 12 s→S). Mann Whitney test (U=19.50, *P*=0.0008): c→C vs s→S, *** *P*<0.001.

**c.** Time mice spent immobile in the forced swim assay (n=10 c→C, 10 s→S). Unpaired student test (t=6.815, *P*<0.0001): c→C vs s→S, *** *P*<0.001.

**d-g**. Mother behaviors in postpartum day 13 (PPD13) in the locomotor activity box. Total **(d)** distance travelled**, (e)** vertical counts, and **(f)** stereotypic behavior, measured in 60 minutes, unpaired t-test, *P* > 0.05 for all tests.

**g.** Time spent in the central and peripheral zones by mothers on PPD13, Two way ANOVA, *P* > 0.05.

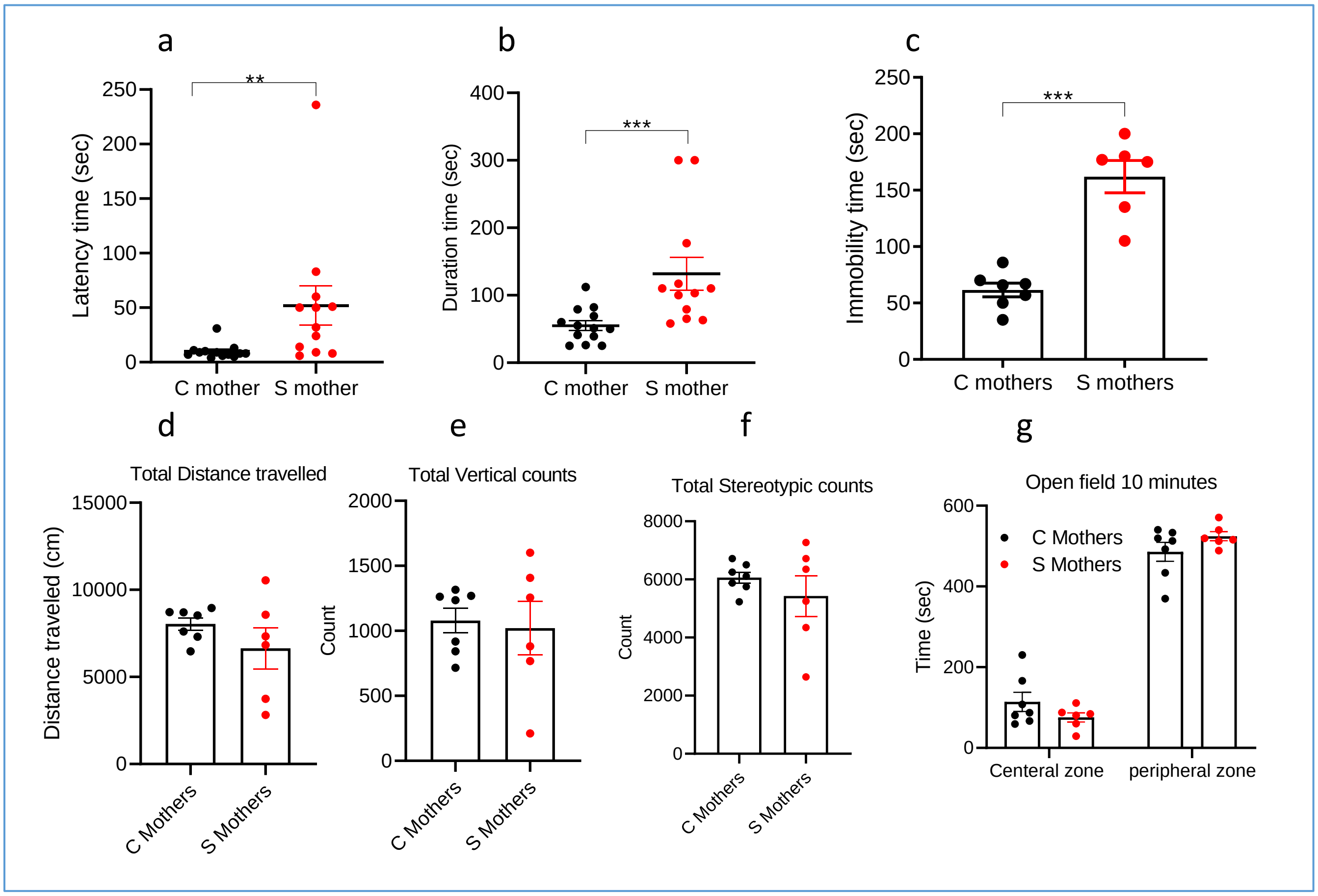

**Figure S2. Bioinformatics analysis of the pathways enriched in DEGs in the brains of neonatal s→S mice**

**a-f:** three top differentially regulated TFs [Fos **(a,b),** Egr1 **(c,d),** and Sp4 **(e,f)**] in the 24-hr mice with their differential target genes identified using **(a,c,e)** MotifMap and **(b,d,f)** ChiPseq

**g,h.** network of differentially regulated TFs **(n)** and RBPs **(o)** in the 24-hr mice, with their differential target genes identified by MotifMap.

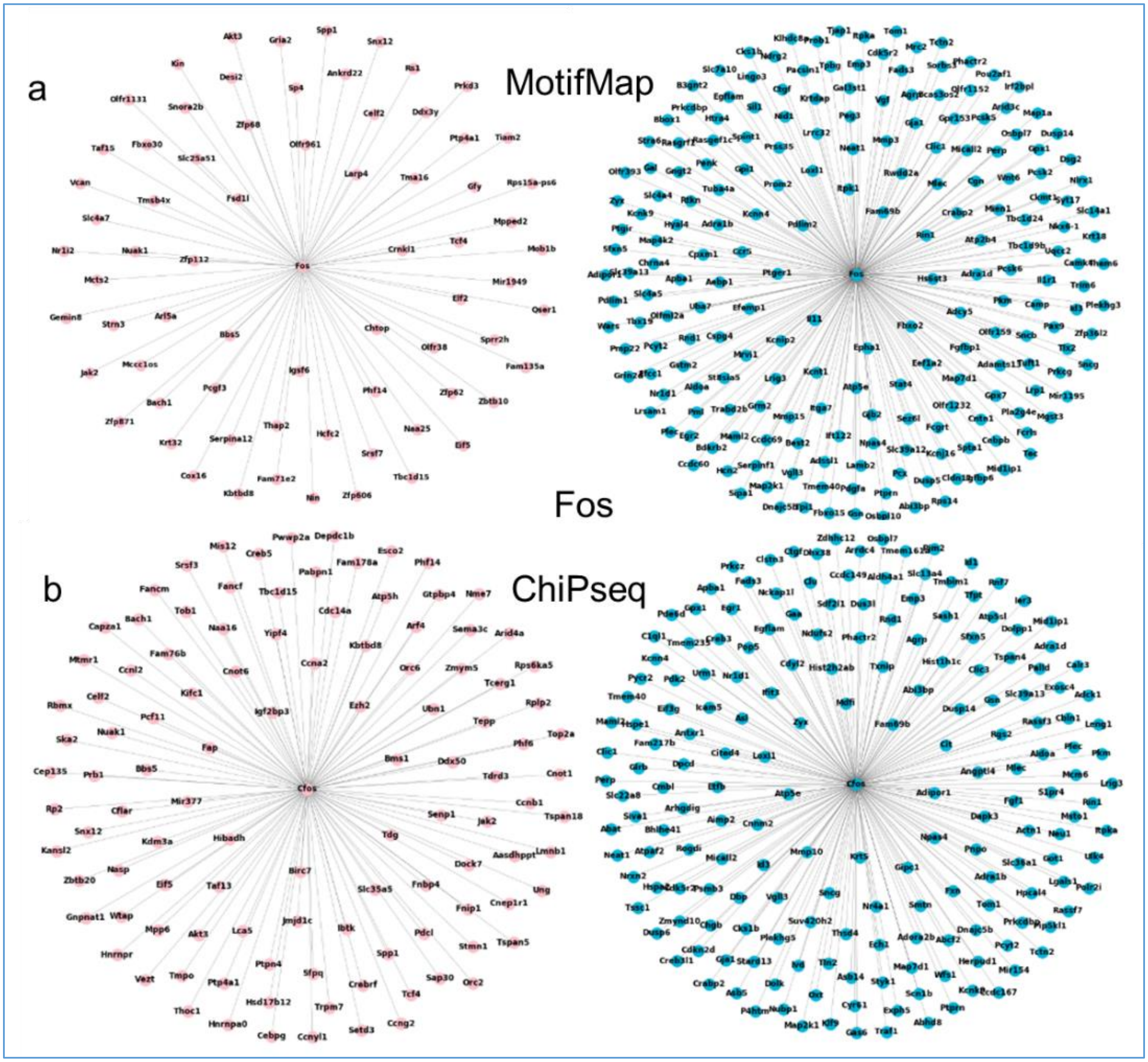

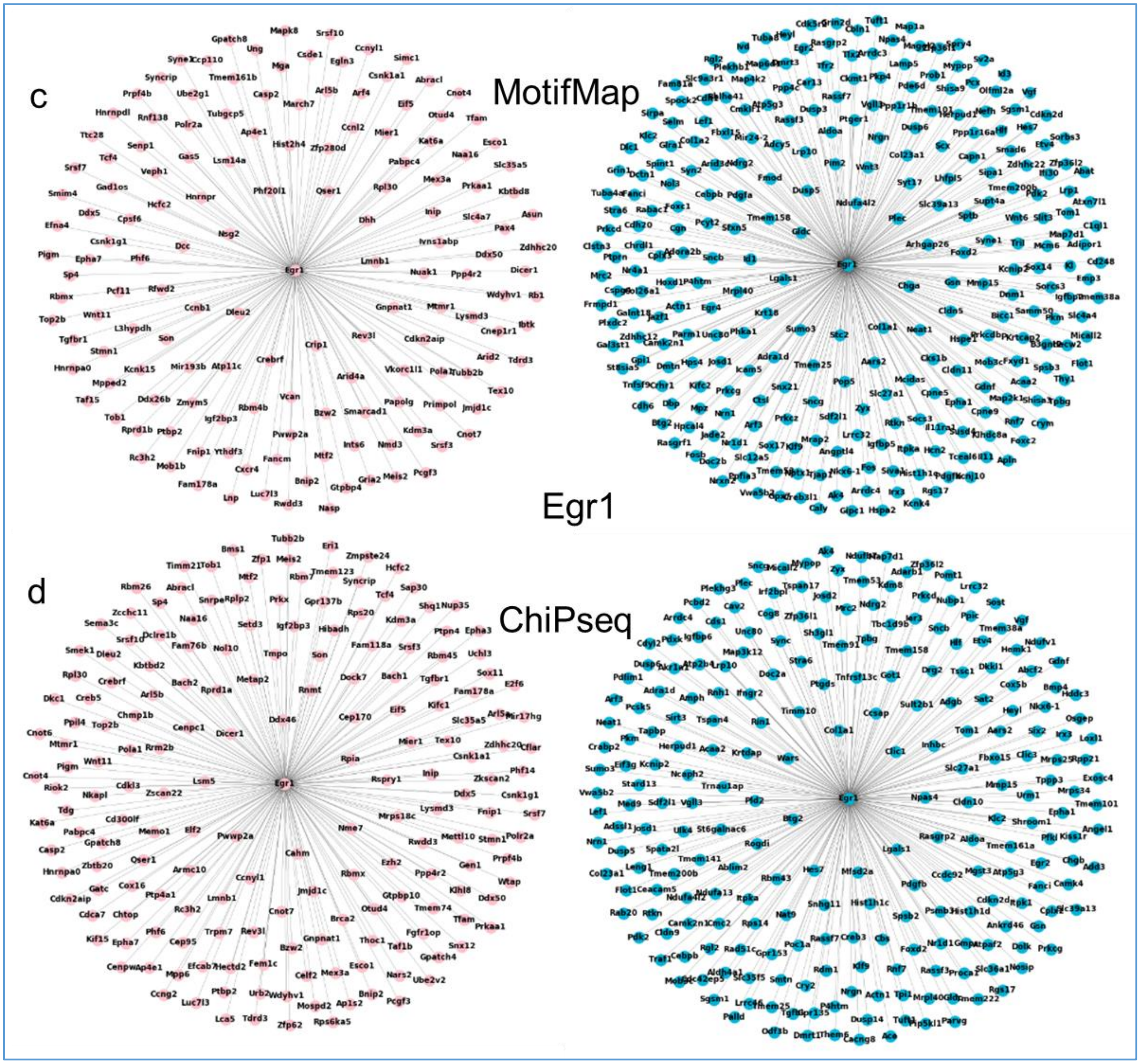
c
MotifMap
Egr1
d
ChiPseq

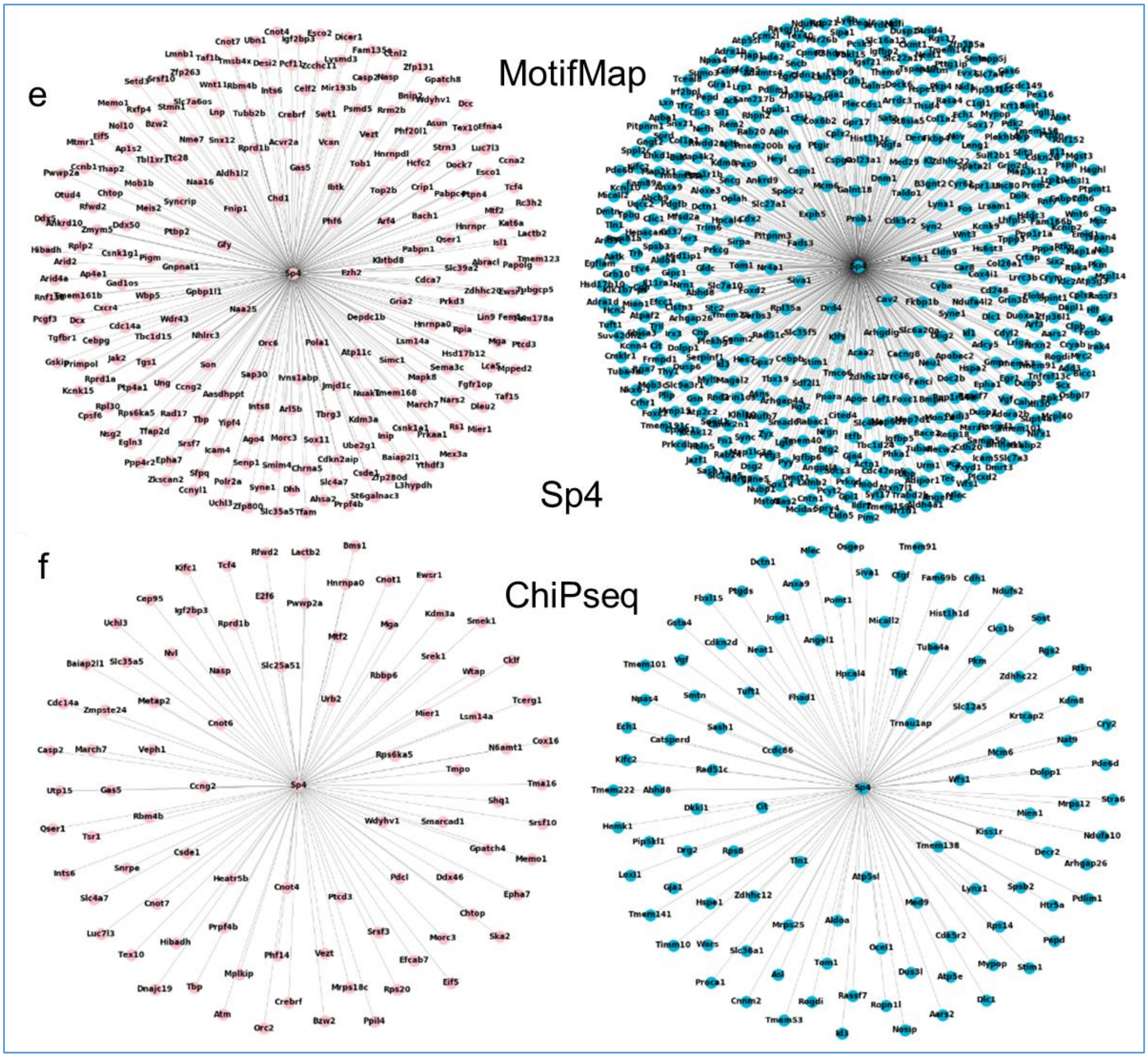
e
MotifMap
Sp4
f
ChiPseq

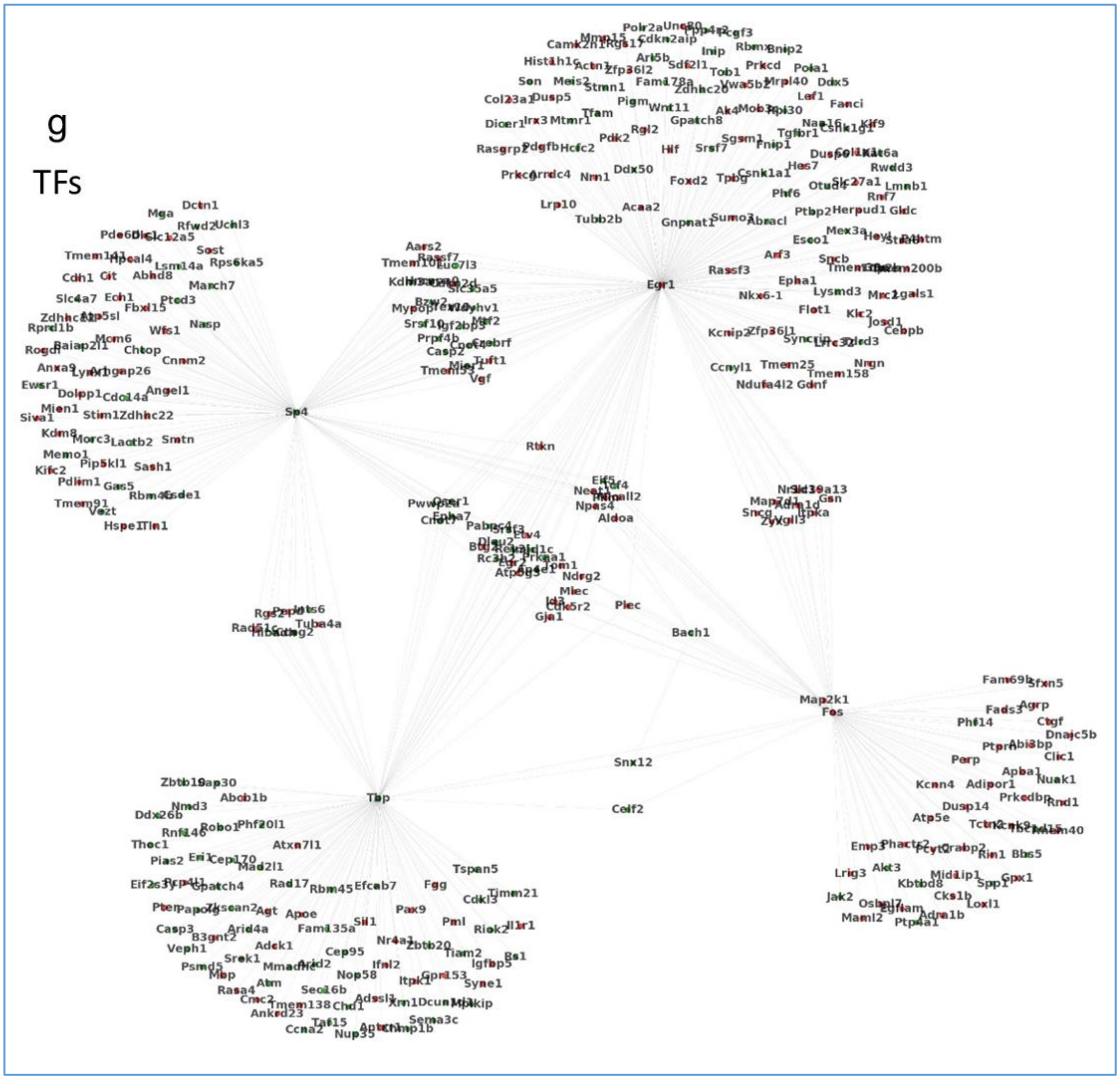
g
TFs
Egr1
Sp4
Map2k1
Fos
Tbp
Rtkn
Snx12
Celf2
Bach1
Plec

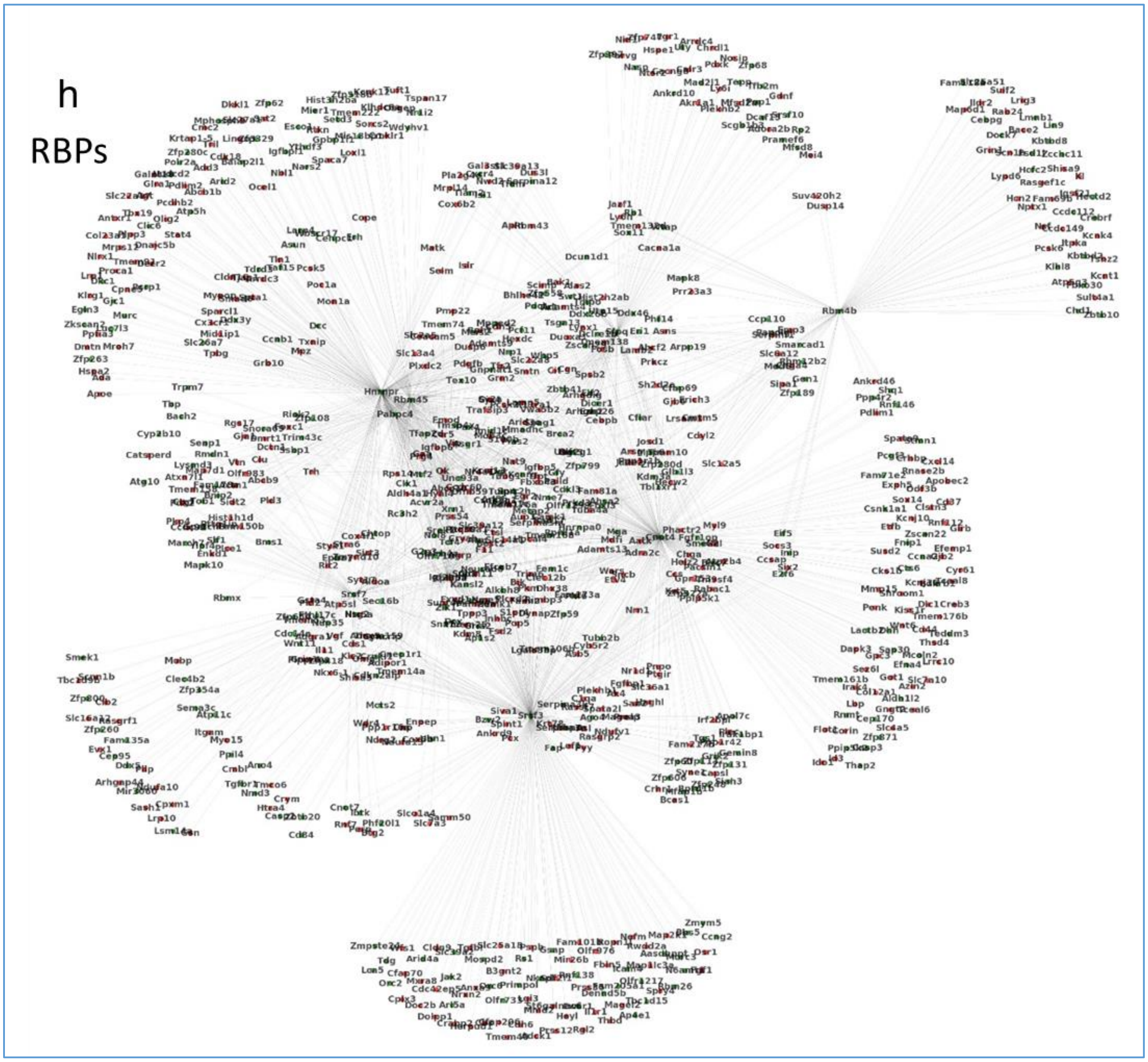
h
RBPs

**Figure S3. Bioinformatics analysis of the pathways enriched in DEGs in the brains of adult s→S mice**

**a-f:** three top differentially regulated TFs [Fos **(a,b),** Egr1 **(c,d),** and Sp4 **(e,f)**] in the adult mice with their differential target genes identified using **(a,c,e)** MotifMap and **(b,d,f)** ChiPseq

**g,h.** network of differentially regulated TFs **(g)** and RBPs **(h)** in the adult mice, with their differential target genes identified by MotifMap.

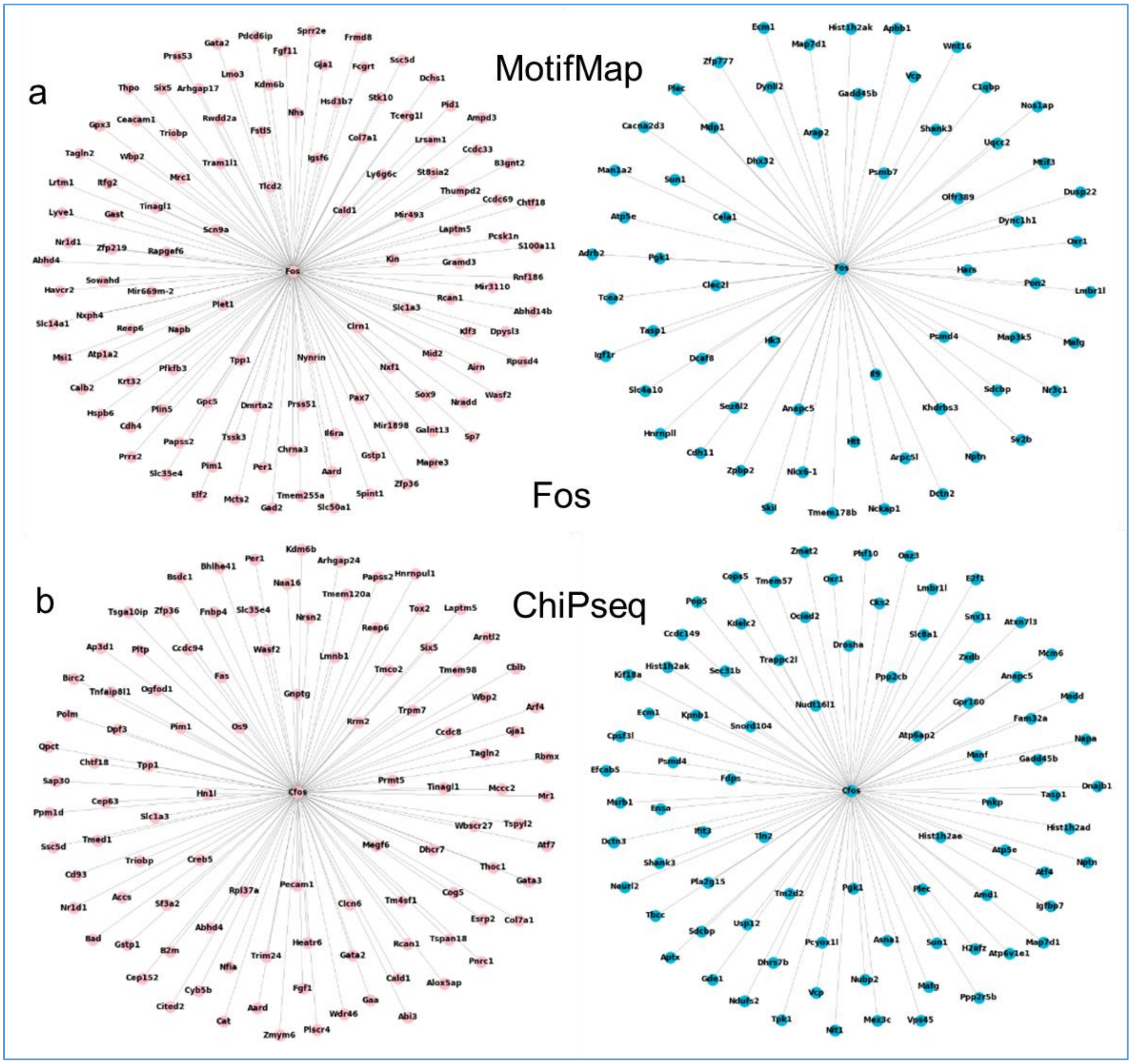

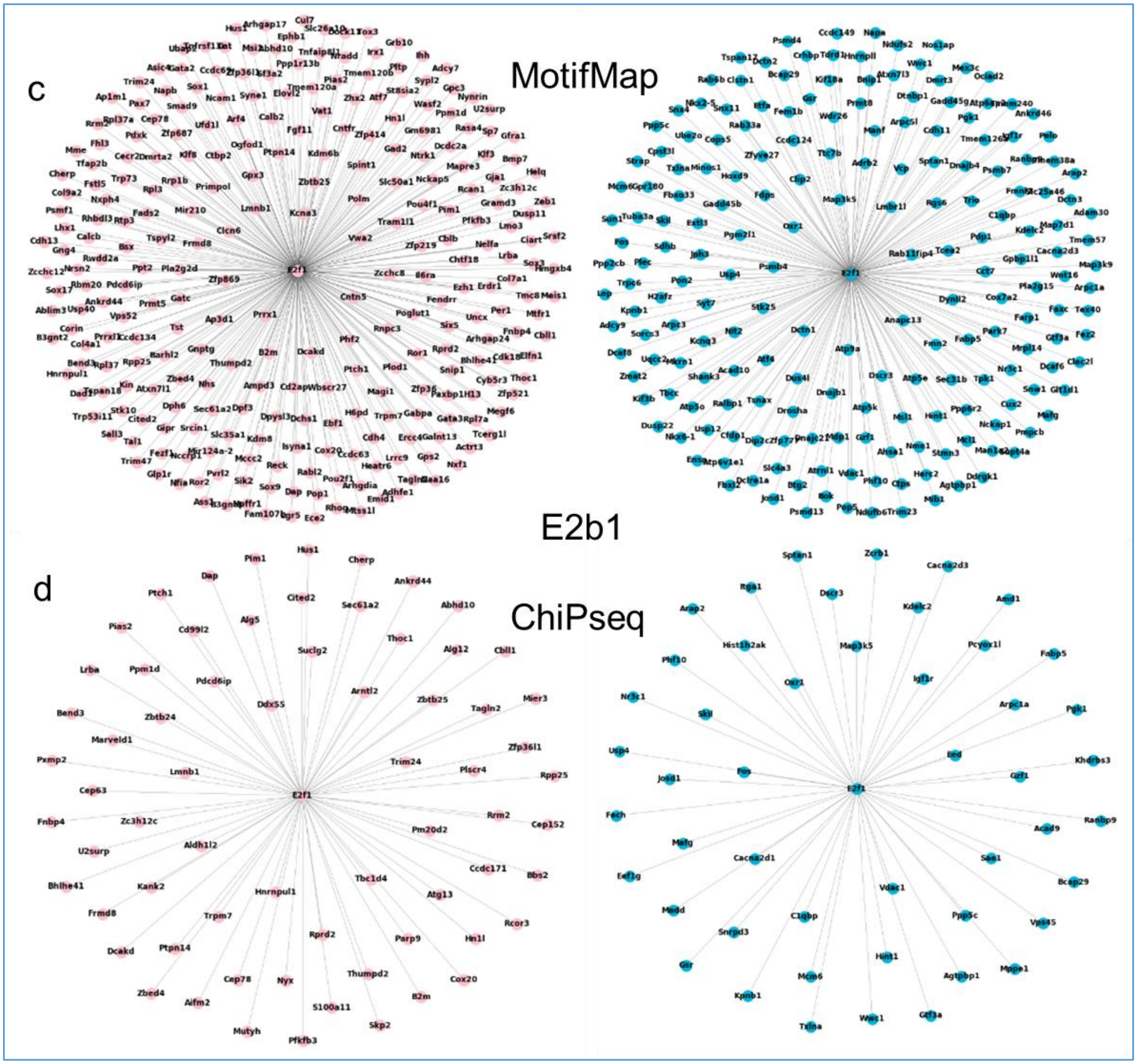
c
MotifMap
E2b1
d
ChiPseq

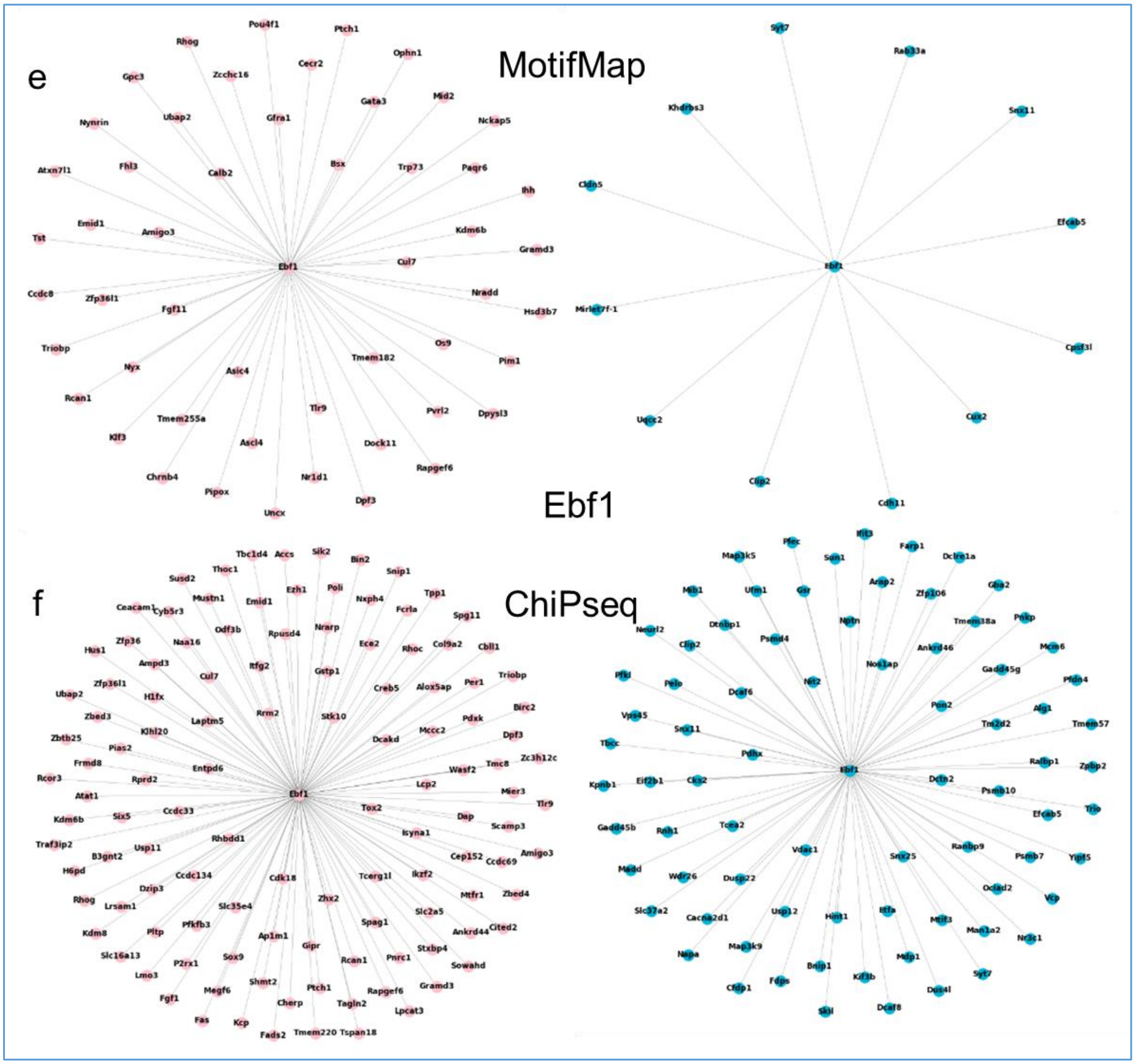
e
MotifMap
Ebf1
f
ChiPseq

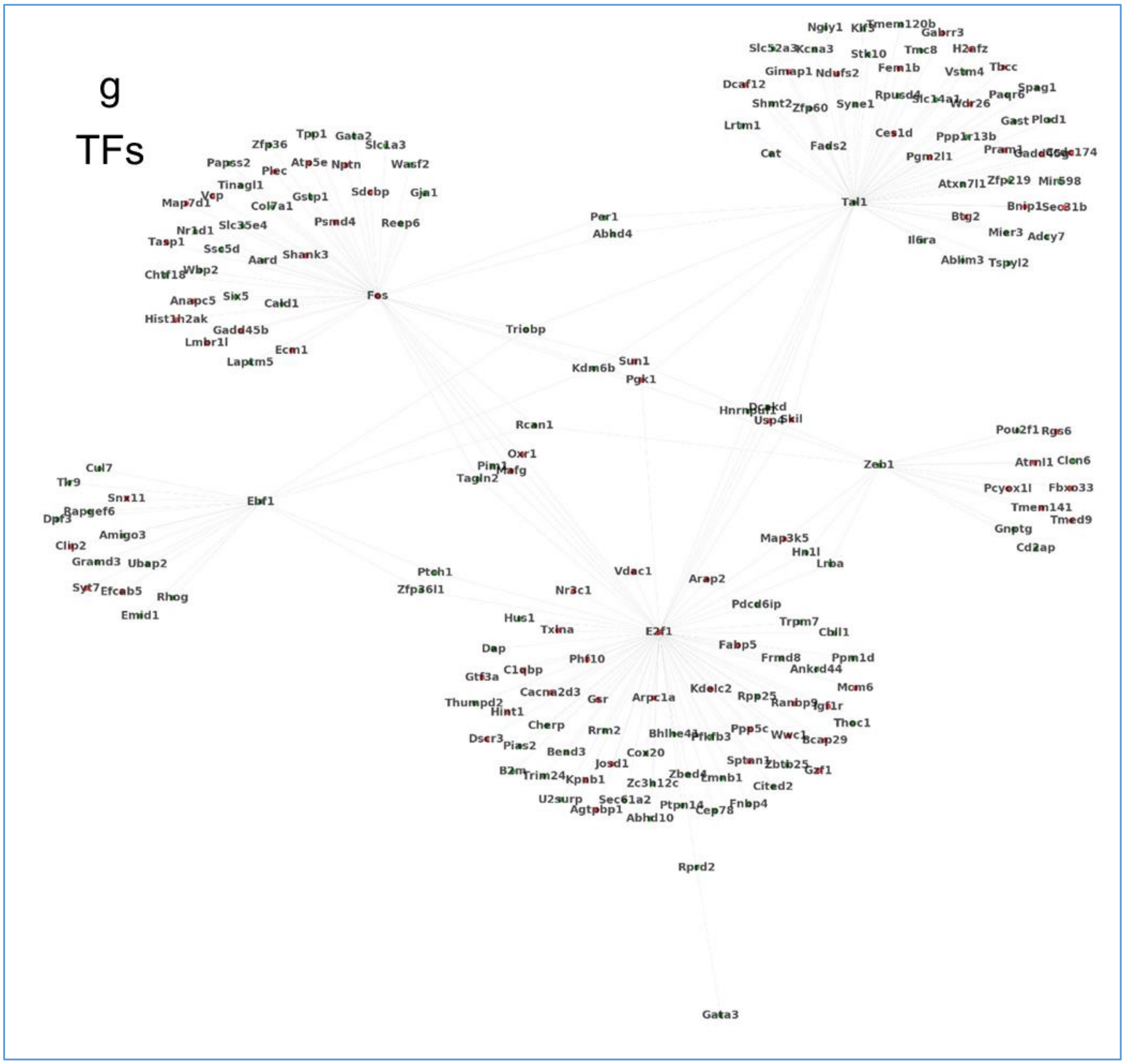
g
TFs
Fos
Tal1
Ebf1
Zeb1
E2f1
Per1
Abhd4
Triobp
Kdm6b
Sun1
Pgk1
Rcan1
Oxr1
Tagln2
Usp4
Skil
Ptch1
Zfp36l1
Vdac1
Nr3c1
Arap2
Gata3
Rprd2
Zfp36
Tpp1
Gata2
Slc1a3
Papss2
Plec
Atp5e
Nptn
Wasf2
Tinagl1
Gstp1
Sdcbp
Gja1
Col7a1
Psmd4
Reep6
Slc35e4
Tasp1
Sse5d
Aard
Shank3
Wbp2
Six5
Caid1
Anapc5
Hist1h2ak
Gadd45b
Lmbr1l
Ecm1
Laptm5
Ngly1
Klf3
Tmem120b
Gabrr3
Slc52a3
Kcna3
Stk10
Tmc8
H2afz
Gimap1
Ndufs2
Fem1b
Vstm4
Tbcc
Dcaf12
Rpusd4
Spag1
Shmt2
Zfp60
Syne1
Wdr26
Paqr6
Lrtm1
Gast
Plod1
Ces1d
Ppp1r13b
Cat
Fads2
Pgm2l1
Pram1
Atxn7l1
Zfp219
Mir698
Bnip1
Sec31b
Btg2
Mier3
Adcy7
Il6ra
Ablim3
Tspyl2
Pou2f1
Rgs6
Atrnl1
Clcn6
Pcyox1l
Fbxo33
Tmem141
Tmed9
Gnptg
Cd2ap
Cul7
Tlr9
Snx11
Rapgef6
Dpf3
Amigo3
Clip2
Gramd3
Ubap2
Syt7
Efcab5
Rhog
Emid1
Map3k5
Hn1l
Lnba
Pdcd6ip
Trpm7
Cbll1
Hus1
Txina
Dap
Fabp5
Frmd8
Ppm1d
Ankrd44
Phf10
C1qbp
Gtf3a
Mcm6
Kdelc2
Cacna2d3
Gsr
Arpc1a
Rpp25
Ranbp9
Igf1r
Thumpd2
Hint1
Thoc1
Cherp
Rrm2
Ppp5c
Wwc1
Bcap29
Dscr3
Pias2
Bend3
Cox20
Sptan1
Gzf1
Josd1
B2m
Trim24
Kpnb1
Zc3h12c
Lmnb1
Cited2
U2surp
Sec61a2
Agtpbp1
Ptpn14
Abhd10
Cep78
Fnbp4

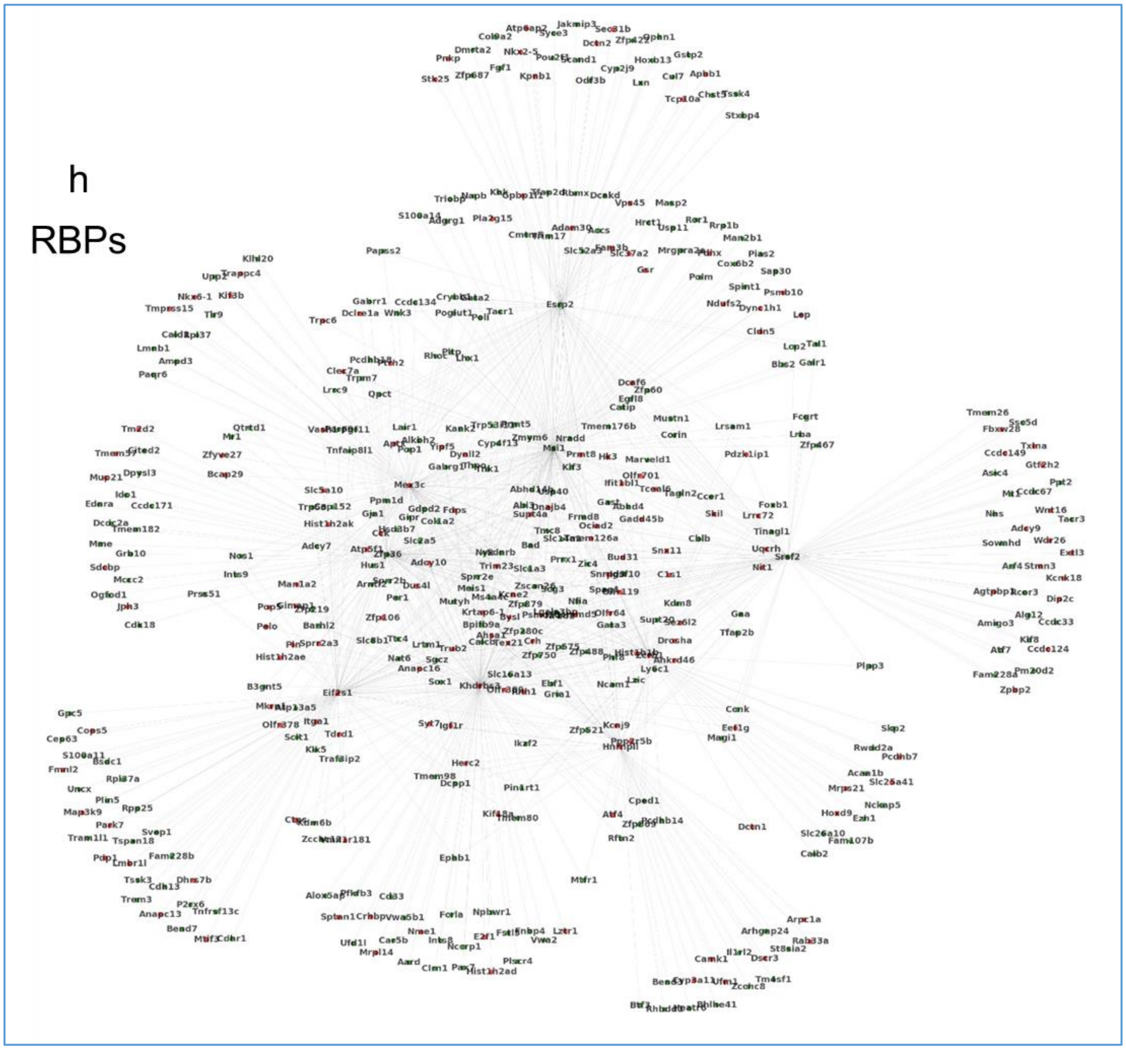
h
RBPs

**Figure S4. Immunohistochemistry analysis of Arc in the brains of adult s→S and c→C mice.** **a,c,e,g,** Representative images of Arc-immunoreactivity in the **(a)** PFC, **(c)** Hip, **(e)** NAc, **(g)** Str **b,d,f,h,** quantification of Arc immuno-positive cells in the **(b)** PFC, **(d)** Hip, **(f)** NAc, (**h)** Str unpaired t-test, PFC: t = 0.198, *P* = 0.85, n = 4; Hip: t=6.678, *P* = 0.001, n = 3 c→C 4 s→S; Nac: t=0.26, *P* = 0.8, n = 4 ; Str: t=2.87, *P* = 0.028, n = 4; Scale bar = 20uM. Values represent mean ± SEM. Frontal cortex (PFC), hippocampus (Hip), nucleus accumbens (NAc), striatum (Str)

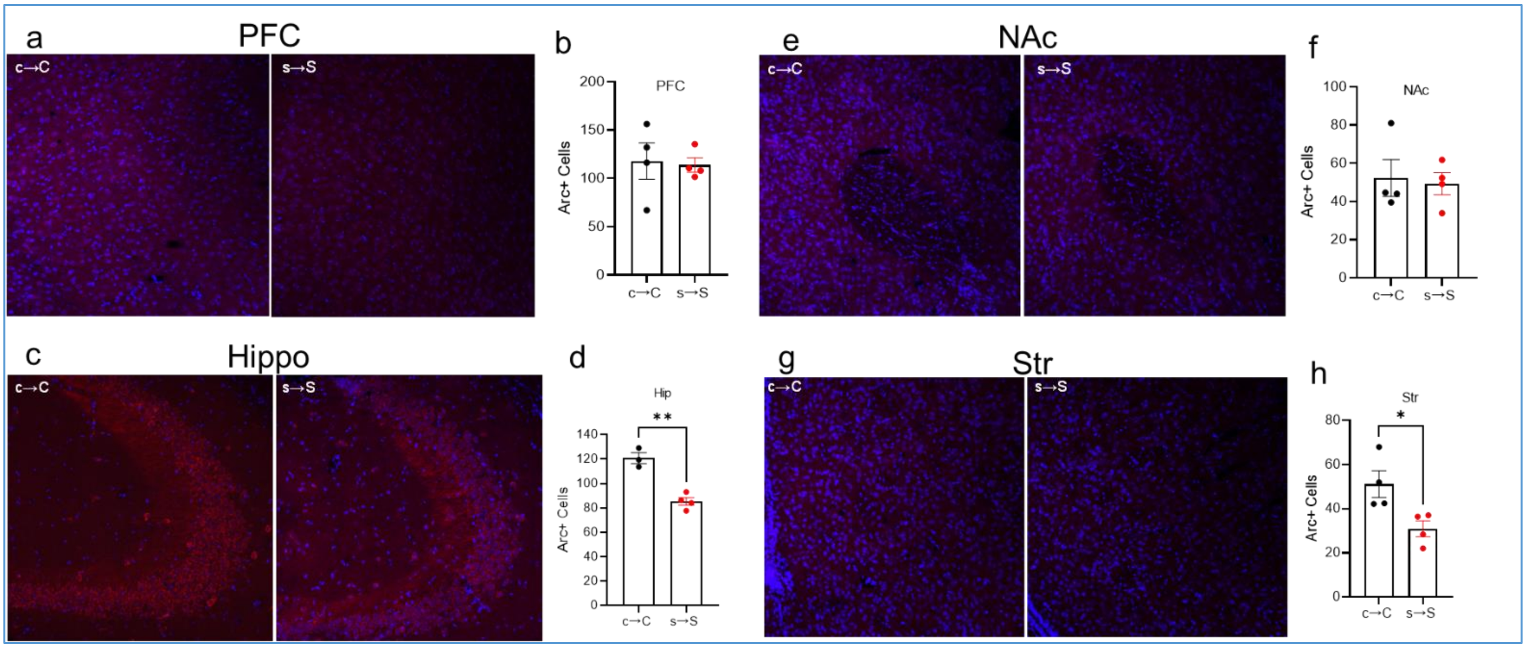

1. Vawter, M.P., et al., *Melanin Concentrating Hormone Signaling Deficits in Schizophrenia: Association With Memory and Social Impairments and Abnormal Sensorimotor Gating.* Int J Neuropsychopharmacol, 2020. **23**(1): p. 53-65.
2. Sanathara, N.M., et al., *Melanin concentrating hormone modulates oxytocin-mediated marble burying.* Neuropharmacology, 2018. **128**: p. 22-32.
3. Alachkar, A., et al., *Prenatal one-carbon metabolism dysregulation programs schizophrenia-like deficits.* Mol Psychiatry, 2018. **23**(2): p. 282-294.
4. Phan, J., et al., *Mating and parenting experiences sculpture mood-modulating effects of oxytocin-MCH signaling.* Sci Rep, 2020. **10**(1): p. 13611.
5. G. Paxinos, K.F., *The Mouse Brain in Stereotaxic Coordinates*, 2001, Academic Press.
6. Schneider, C.A., W.S. Rasband, and K.W. Eliceiri, *NIH Image to ImageJ: 25 years of image analysis.* Nat Methods, 2012. **9**(7): p. 671-5.
7. Kayala, M.A. and P. Baldi, *Cyber-T web server: differential analysis of high-throughput data.* Nucleic Acids Res, 2012. **40**(Web Server issue): p. W553-9.
8. Baldi, P. and A.D. Long, *A Bayesian framework for the analysis of microarray expression data: regularized t -test and statistical inferences of gene changes.* Bioinformatics, 2001. **17**(6): p. 509-19.
9. Cerami, E.G., et al., *Pathway Commons, a web resource for biological pathway data.* Nucleic Acids Res, 2011. **39**(Database issue): p. D685-90.
10. Kamburov, A., et al., *ConsensusPathDB--a database for integrating human functional interaction networks.* Nucleic Acids Res, 2009. **37**(Database issue): p. D623-8.
11. Liu, Y., et al., *MotifMap-RNA: a genome-wide map of RBP binding sites.* Bioinformatics, 2017. **33**(13): p. 2029-2031.
12. Daily, K., et al., *MotifMap: integrative genome-wide maps of regulatory motif sites for model species.* BMC Bioinformatics, 2011. **12**: p. 495.
13. Consortium, E.P., *An integrated encyclopedia of DNA elements in the human genome.* Nature, 2012. **489**(7414): p. 57-74.
